# Seeing the chemistry of biomolecular condensates: *in situ* mapping of composition and water content

**DOI:** 10.1101/2025.10.20.683555

**Authors:** E. Sabri, A. Mangiarotti, C.N.Z. Schmitt, R. Dimova

**Affiliations:** Max Planck Institute of Colloids and Interfaces, Science Park Golm, 14476 Potsdam, Germany; Centro de Investigaciones en Química Biológica de Córdoba (CIQUIBIC), CONICET, X5000HUA Córdoba, Argentina; Departamento de Química Biológica Ranwel Caputto, Facultad de Ciencias Químicas, Universidad Nacional de Córdoba, X5000HUA Córdoba, Argentina

## Abstract

Biomolecular condensates are membraneless cellular organelles that form via liquid–liquid phase separation of proteins and nucleic acids. Their functional roles are tightly coupled to material properties like viscosity and hydrophobicity, which serve as key markers of cellular state. However, conventional determination of condensate composition and water content relies on invasive procedures that can damage samples. Here, we introduce Raman spectroscopy coupled with spectral phasor analysis as an *in situ*, label-free approach to resolve the chemical profiles and molecular concentrations within both the dense and dilute phases of biomolecular condensates. This method outperforms traditional regression and deconvolution approaches, yielding a precise readout of client molecule partitioning. By accounting for contributions of the protein backbone to the Raman spectra of condensates, we assess the signature of “solid-like” hydrogen-bonded water from protein hydration, revealing that most water molecules within condensates retain bulk, “liquid-like” properties. Finally, using environment-sensitive fluorescent probes, we demonstrate that macromolecular structure and water content—rather than hydrogen bonding alone—drive condensate hydrophobicity; notably, the dense phase remains predominantly water-rich even at low apparent dielectric constants.

## 1. INTRODUCTION

Biomolecular condensates form via the self-assembly of proteins and/or nucleic acids, a process known as liquid–liquid phase separation (LLPS). These protein-rich aqueous droplets act in cells as function-specific membraneless organelles that can selectively encapsulate small molecules^1, 2^, modulate the rate of chemical reactions^1, 3^ and participate in gene regulation^4^ to mention a few. Biomolecular condensates can transition from their liquid state to gel-like or solid assemblies with significantly different material properties which act as sensitive markers of their physiological state in health and disease^5-8^. The material properties of condensates are commonly interpreted as emergent properties, governed by the structural features of biopolymers composing the condensate^5, 8-10^.

Beyond the properties of the biopolymers themselves, water plays a dual role in cellular organization. It shapes protein structure^11-14^ and biochemical activity^15^, while hydrated proteins, in turn, alter the dynamics of their surrounding water shells, yielding properties fundamentally distinct from bulk water^9, 15^, and influencing the partitioning of small molecules within condensates^1, 2, 16^. In crowded biomolecular environments, the dynamics of water molecules are entropically penalized^17-20^. LLPS allows cells to reversibly reorganize intracellular water and buffer changes in cytosolic conditions, thereby contributing to cellular homeostasis under stress^9, 18^. While numerical estimates suggest that dynamically constrained water may extend over several hydration layers (up to approximately 6) around proteins in solutions^21-24^, the corresponding volume fractions of constraint versus bulk-like water in condensates remain experimentally undetermined. Consequently, a central and unresolved question is whether the functional properties of condensates emerge primarily from specific molecular features of the phase-separating proteins^2, 25^—such as protein size, the presence of intrinsically disordered regions, or the number of hydrophobic residues—or from altered properties of solvent molecules within condensate droplets compared to the bulk solvent^1, 2^. This question is particularly relevant for understanding the molecular determinants of condensate hydrophobicity, as mounting evidence indicates that hydrophobic properties strongly influence the partitioning efficiency of small client molecules^2, 16, 26^, with important implications for cellular metabolism and drug processing ^27^. Crucially, here we distinguish between mesoscale water content and the local “hydrophobicity” reported by environment-sensitive dyes, which only probe their immediate nanoscale surroundings^17-20^. While water properties and dynamics are altered in the vicinity of proteins, an open question we address in this study is whether dye-reported hydrophobicity directly reflects the actual water fraction within the dense phase of condensates.

Nonetheless, a major limitation of existing approaches is that quantifying the water content within condensates, identifying the molecular constituents of their biopolymer components, or measuring the partitioning efficiency of exogenous small molecules typically requires invasive and destructive steps such as sample drying^28-30^, which are inherently incompatible with *in vivo* measurements. Raman spectroscopy has emerged as a powerful non-invasive approach for correlating microscopic chemical and structural variations with macroscopic biophysical properties in biological and biomimetic systems^31-33^.

Existing analytical strategies generally fall into two broad categories. A first class of approaches relies on qualitative spectral metrics, such as intensity ratios between selected Raman shifts^31, 34-41^ or shifts in local spectral maxima^31, 32, 34, 42^. While effective for detecting relative spectral changes, these metrics only probe restricted subsets of vibrational modes and can remain sensitive to spectral degeneracies, where distinct physicochemical processes generate similar spectral signatures^42^.

Alternatively, quantitative compositional analyses typically attempt to deconvolve the respective contributions of multiple molecular constituents to the measured spectrum. This is commonly achieved either through fitting procedures, such as Gaussian decomposition^34, 43-46^ or multivariate curve resolution by alternating least-squares (MCR-ALS)^47-49^, or through regression-based frameworks including principal component regression (PCR)^50, 51^ and partial least-squares regression (PLSR)^52, 53^. Although highly powerful, these approaches generally require either prescribing a priori the number of spectral degrees of freedom, defining explicit fitting models, or constructing extensive calibration datasets to relate spectral features to physicochemical variables.

Here, we present a combination of Raman spectroscopy and spectral phasor analysis that provides a fit-free and minimally supervised framework for quantifying the respective volume fractions of polymers and water inside biomolecular condensates. By exploiting the linear geometry of spectral mixtures in Fourier space, this approach enables direct compositional mapping using minimal a priori information regarding the system composition. This approach enables the assessment of small-molecules partitioning within condensates while simultaneously quantifying the fraction of hydrogen-bonded water molecules hydrating the polymer matrix. We show that these measurements are robust irrespective of molecular size, of the type of LLPS (i.e. associative or segregative), or macromolecular structural complexity, providing a generalizable benchmark for single-droplet condensate compositional analysis. Building on this framework, we quantitatively investigate the structural properties of water molecules across all tested condensates. Finally, we use ACDAN as an environment-sensitive fluorescent reporter of condensate hydrophobicity^54^ to examine how water content and biopolymer structural motifs correlate with the hydrophobic properties of biomolecular condensates.

## 2. RESULTS

### 2.1 Raman spectroscopy quantifies hydrogen-bonded atoms in organic compounds

Raman spectroscopy relies on inelastic photon scattering to extract information about the abundance of different types of chemical bonds in a sample, based on the vibrational modes of the bonded atoms. This technique has been widely used in biology as a non-invasive probe of subtle changes in intermolecular interactions, bond geometry, and conformational rearrangements at the microscopic scale^31, 32, 43, 55^. From a theoretical perspective, vibrational stretching modes of interatomic bonds can be approximated by those of a harmonic oscillator, as presented in Fig. 1a. Such modes are defined primarily by two parameters: the bond strength modeled by a spring constant *k*_*bond*_, and the effective mass of the bonded atoms 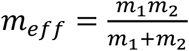, where *m*_1_ and *m*_2_ are the masses of the different atoms. At room temperature, thermal energy (*k*_*b*_*T* ≈ 4.11 × 10^−21^J) is typically one order of magnitude lower than the energy gap that separates the vibrational ground state from the first excited vibrational state in organic molecules^31, 32^. As a result, vibrational modes are not thermally populated under ambient conditions.

**Figure 1:**
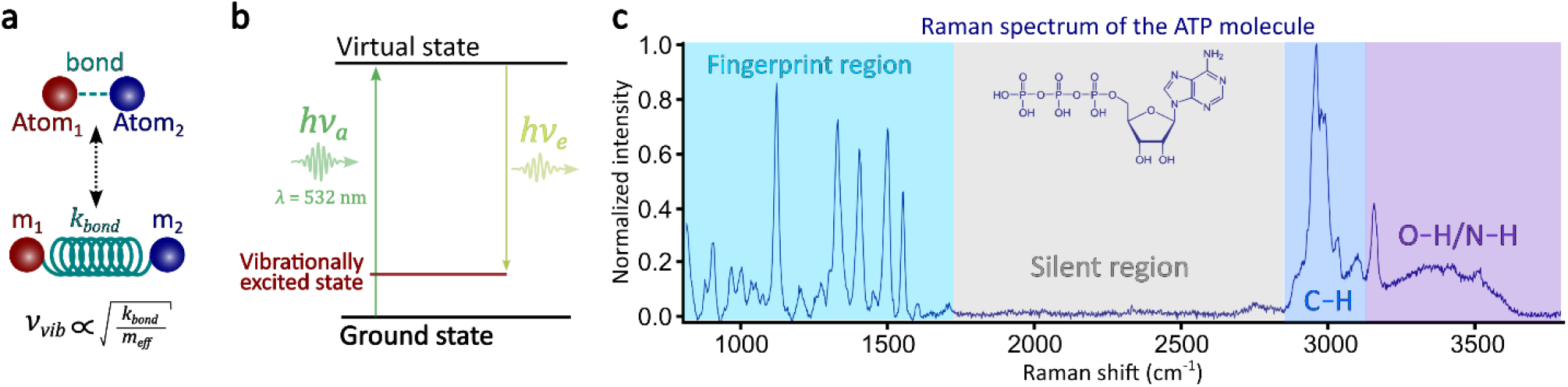
Physical principles underlying Raman vibrational spectroscopy. **(a)** Schematic representation of an interatomic bond modeled as a harmonic oscillator, in which two atoms of masses *m*_1_ and *m*_2_ are connected by a spring with force constant *kbond*, and of which the vibrational modes of frequency *νvib* scale with 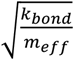. **(b)** Jablonski diagram illustrating the energy transitions involved in Stokes Raman scattering. Absorption of an incident photon promotes the system to a virtual excited state, followed by emission of a lower-energy photon. The red and black levels represent the vibrationally excited and ground states, respectively. **(c)** Normalized Raman spectrum of pure (lyophilized) ATP plotted as a function of Raman shift 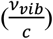. The spectral regions associated with C–H stretching vibrations (2800-3100cm^-1^) and O–H/N–H stretching vibrations (3100-3800cm^-1^) are highlighted in blue and purple, respectively^31^. The fingerprint region^33^ is highlighted in cyan, while the silent region^33^ is shown in grey.

However, these vibrational modes can be excited by exposing the sample to a monochromatic light source, resulting in a cascade of photon absorption-emission events that yield information about the energy gap separating these modes^55^ (Fig. 1b). In the context of Stokes Raman scattering, the absorption of a photon promotes the system to a virtual excited state, followed by the emission of a lower-energy photon. The difference in energy *h*(*ν*_*a*_ − *ν*_*e*_) between the absorbed and emitted photons – the Stokes Raman shift – matches the energy of a specific molecular vibration *hν*_*vib*_ which defines the final vibrational state of the molecule.

Figure 1c presents the Raman spectrum of adenosine triphosphate (ATP) as an illustrative example of the spectral profile of a canonical organic compound. The spectrum distinguishes between spectral characteristic regions associated with specific chemical features. The low-frequency fingerprint region (400–1800 cm^-1^)^33^ is dominated by vibrations involving heavier atoms linked by single or double covalent bonds, such as C–C (≈800 cm^-1^) and C=C (≈1600 cm^-1^), whereas vibrational modes of more strongly bonded counterparts, such as C≡C (≈2160 cm^-1^) or C≡N (≈2230 cm^-1^), appear in the silent region (Fig. 1c)^33^. By contrast, interatomic vibrational modes involving lighter atom pairs that include hydrogen appear at higher frequencies and are of particular relevance here, namely 2700-3100 cm^-1^ for C–H stretching^31^, 3100-3800 cm^-1^ for O–H stretching^31^ and 3100-3400 cm^-1^ for N–H stretching^56^.

### 2.2 Raman intensities scale linearly with interatomic bond abundance

To exploit the full spectral information contained in the measured Raman signals, we first establish the correspondence between the Raman intensity integrated over a given spectral region and the volume fraction of the corresponding chemical bonds within the sample. Figure 2 presents a proof-of-principle demonstrating that the relative abundance of C–H, O–H, and N–H bonds can be directly quantified from integrated Raman intensities over their respective vibrational regions.

**Figure 2:**
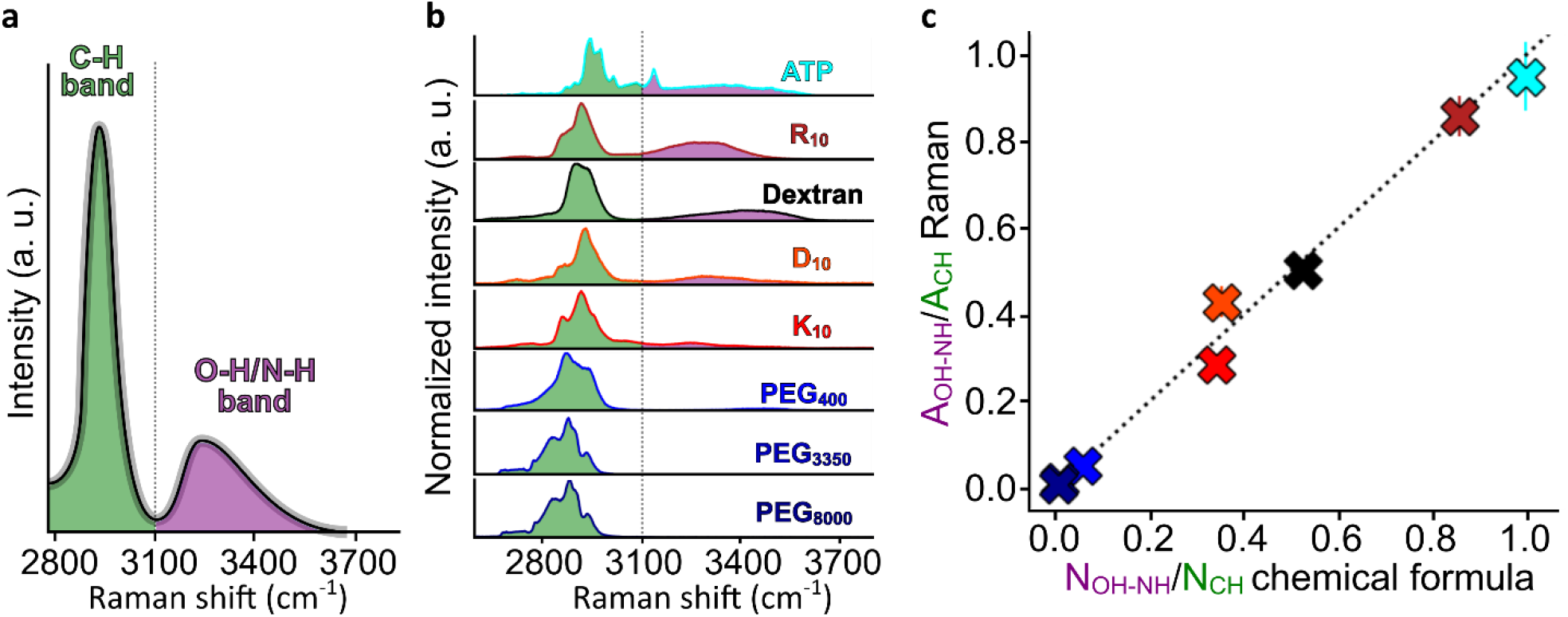
Raman spectroscopy enables quantitative assessment of interatomic bond abundance in polymers. **(a)** Schematic illustration of the typical Raman spectrum of a molecule used in this study. The areas under the curve corresponding to C–H stretching vibrations (*ACH*, green) and to O–H/N–H stretching vibrations (*AOH*−*NH*, purple) are highlighted. **(b)** Normalized Raman spectra for different molecules (in lyophilized form) with known chemical compositions displayed over the 2700–3800 cm^−1^ spectral region and ordered by decreasing O–H/N–H bond abundance from top to bottom. Spectral profiles consist of five independent measurements that were normalized and averaged for each molecule. **(c)** Linear correspondence between spectrally inferred and stoichiometrically calculated bond ratios. The ratio *AOH*−*NH*/*ACH* obtained from integrated Raman intensities is plotted against the corresponding ratio *NOH*−*NH* /*NCH* of the known number of C–H and sum of O–H/N–H bonds calculated from the known chemical formulas of the molecules shown in (b) in corresponding color.

Figure 2a illustrates the typical Raman spectral profile of a protein, with C–H stretching modes quantified by the area under the curve in the 2700–3100 cm^−1^ region^31^, denoted *A*_*CH*_, whereas O–H and N–H stretching modes are quantified by the area under the curve in the 3100–3800 cm^−1^ region^31, 56^ denoted *A*_*OH*−*NH*_. The chemical formulas of all compounds are known *a priori*. We denote the number of C–H bonds as *N*_*CH*_ and the combined number of O–H and N–H bonds as *N*_*OH*−*NH*_. Figure 2b presents the normalized Raman spectra of the polymers and biomolecules considered in this study over the 2700–3800 cm^−1^ region, ordered by decreasing O–H/N–H bond abundance from top to bottom. As a result, for several pure protein compounds, the intrinsic O– H/N–H Raman signal overlaps with the spectral region commonly associated with bulk water vibrations^36, 44^. This overlap is particularly important when analyzing protein-rich aqueous environments, where intensity variations over the 3100–3800 cm^−1^ region are often interpreted as changes in water dynamics – through enhanced or weakened hydrogen bonding – resulting from protein hydration^44, 57^. However, the data presented in Fig. 2b suggests an alternative interpretation, whereby such spectral variations may partially originate from the intrinsic chemical composition of the protein backbone itself. Lastly, Fig. 2c compares the relative abundance of C–H versus O– H/N–H bonds inferred from the ratio of integrated Raman intensities (vertical axis) with the corresponding ratios calculated from the known chemical compositions of the molecules (horizontal axis). These results reveal a clear identity relation between the stoichiometric bond ratios and their corresponding integrated spectral intensities. Taken together, these results demonstrate that, for composite Raman spectra, the molecular chemical signatures encoded in the relative abundance of distinct classes of chemical bonds can be reliably disentangled using a Fourier-space analysis.

### 2.3 Raman spectral phasor analysis quantitatively resolves polymer and water compositions in crowded solutions and phase-separated systems

Over the past few years, Raman spectroscopy has gained momentum in the study of biomolecular condensates^58^, with a particular emphasis on identifying LLPS propensity markers encoded in protein structure^34, 59^, resolving cellular signatures of LLPS-mediated stress responses^32, 36^, and more recently, probing the role of water dynamics in shaping the macroscopic material properties of condensates^35, 36, 44, 57, 60^. In this latter context, special attention has been given to changes in the intensity profile of the O–H bond stretching band (3100-3800 cm^-1^), where altered spectral profiles are commonly interpreted as variations in hydration levels and water dynamics relative to protein or nucleic acid content through hydrogen bonding^35, 44, 57, 60^. However, such interpretations implicitly assume that contributions from the O–H and N–H bonds in the polymer backbone are insignificant over this spectral region, an assumption that is not generally valid in crowded or phase-separated environments as demonstrated in Fig. 2c.

To deconvolve the contributions of O–H and N–H bonds in the protein backbone of macromolecules from the intrinsic O–H profile of water, we leverage the Fourier-transform properties of Raman spectra. Our aim is to explore whether spectral phasor analysis can be used to directly quantify the respective volume fractions of water and proteins or polymers in phase-separated systems. We first consider aqueous binary mixtures of polyethylene glycol (PEG), a well-characterized and affordable water-soluble polymer system. Raman spectra acquired over the 2700-3800 cm^-1^ region for PEG_400_-water mixtures of varied polymer content are presented in Fig. 3a.

**Figure 3:**
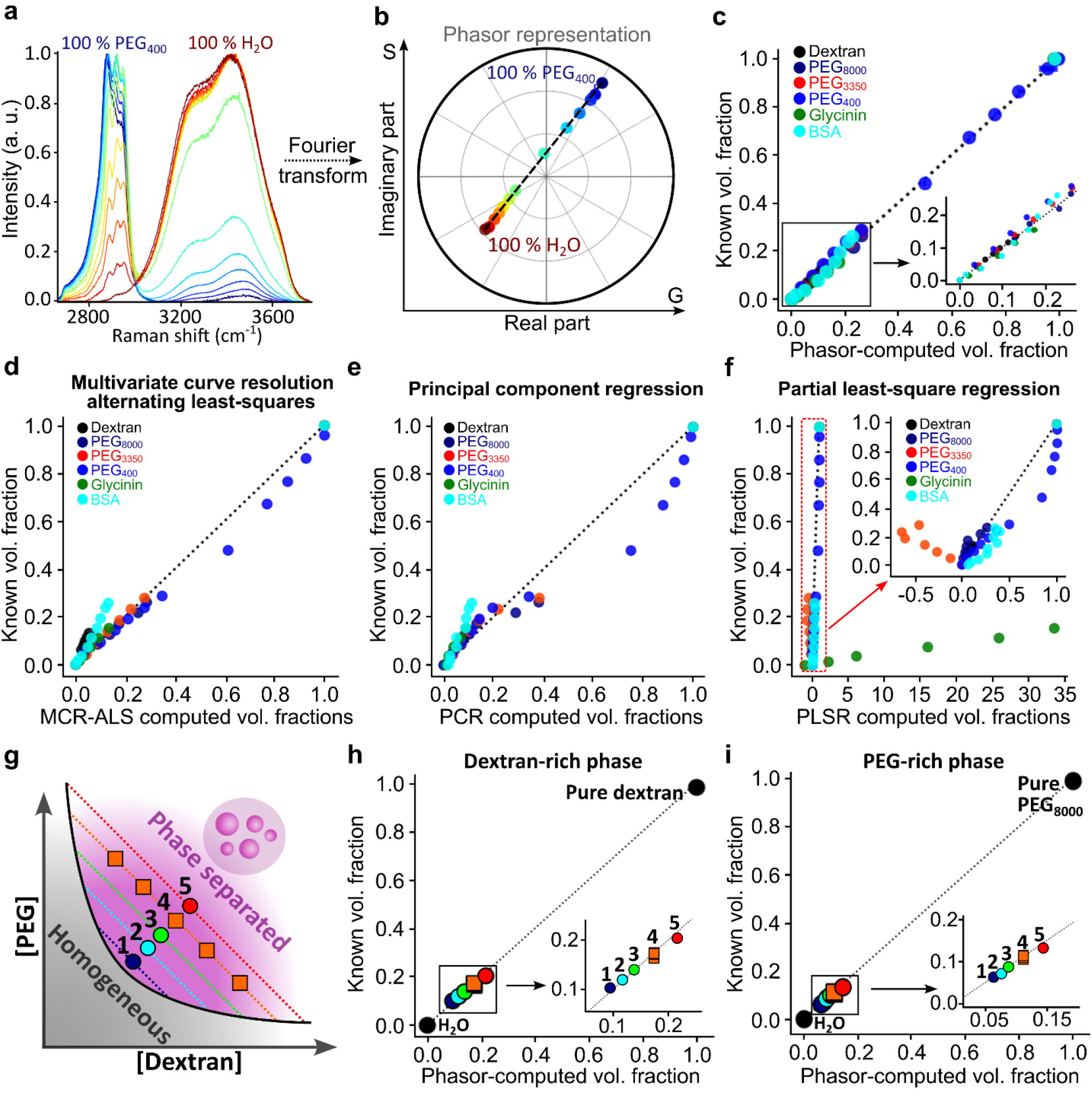
Spectral phasor analysis quantitatively resolves the volume fraction occupied by macromolecules in homogeneous aqueous solutions and phase separated systems. **(a)** Normalized Raman spectra of PEG400-water mixtures with polymer volume fractions of 100%, 92.7%, 83.5%, 74.2%, 64.9%, 46.4%, 27.8%, 23.2%, 18.5%, 13.9%, 9.3%, 4.6% and 0% PEG400 (from blue to red, n=3 for each condition). **(b)** Corresponding spectral phasor representation obtained from phasor representation of the Fourier transform of the spectra in (a). The black dashed line is a guide to the eye connecting the 100% H2O and 100% PEG400 reference points. **(c)** Comparison between macromolecule volume fractions computed from phasor analysis (horizontal axis, Eq. M8) and experimentally controlled macromolecule volume fractions in aqueous solutions (vertical axis). The inset shows a magnified view. The black dotted identity line is a guide to the eye confirming that the phasor approach systematically provides an accurate readout of polymer-to-water ratio. **(d-f)** Same as in (c) for MCR-ALS, PCR and PLSR respectively. The inset in (f) shows a magnified view. **(g)** Schematic phase diagram of a PEG-dextran ATPS^62^. The purple and dark grey regions respectively indicate phase-separated and homogeneous regimes; the solid black curve is the binodal. Colored dotted lines represent tie lines whose intersections with the binodal define the compositions of the PEG-rich and dextran-rich phases. Colors and symbol shapes schematically correspond to the data points shown in panels (h) and (i). Symbols of identical color (orange) correspond to measurements performed along the same tie line, for which the compositions of the two coexisting phases are fixed, while only their relative volume fractions vary. **(h, i)** Polymer volume fractions of dextran in the dextran-rich phase (h) and PEG8000 in the PEG-rich phase (i) obtained from phasor analysis (horizontal axes) compared with values inferred from the experimental phase diagram reported in ref. ^62^ (vertical axes). The insets show a magnified view. The numbers 1-5 indicate distinct tie lines as shown in panel (g) where three independent measurements were performed for each position represented on the phase diagram.

Figure 3b shows the corresponding spectral phasor representation (see Methods and Fig. S1), where the horizontal (G) and vertical (S) coordinates correspond to the real and imaginary parts of the first harmonic component of the Fourier transform of the Raman spectrum^61^. In phasor space, the spectra of all PEG_400_-water mixtures (represented as points) align along the straight line connecting the pure water and pure PEG_400_ reference points, consistent with the linearity of phasor-space geometric operations for binary mixtures^61^. Figures S2, S3 demonstrate that this behavior generalizes across all tested polymer-water and protein–water systems.

Exploiting this geometric property (Fig. S1 and Methods), we computed the polymer volume fraction *V*_*phasor*_ directly from the phasor coordinates (Eq. M8). Figure 3c compares these phasor-derived values with the known polymer volume fractions *V*_*known*_ used to prepare the solutions. The latter were calculated as 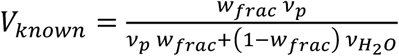, where *w*_*frac*_ is the chosen weight fraction of the polymer in the mixture, *ν*_*p*_ is the specific volume of the polymer, and 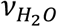 the specific volume of water, all listed in Table S1. The excellent agreement between *V*_*phasor*_ and *V*_*known*_ across a broad range of polymers demonstrates that the spectral phasor provides a robust and quantitative readout of polymer-to-water composition in crowded aqueous solutions.

Importantly, the sensitivity of the phasor approach scales inversely with the total width of the analysed spectral window. This is relevant for mitigating localized spectral distortions, such as peak attenuations and frequency shifts associated with Fermi resonance^32, 43, 63^. In practice, such effects can be reduced by broadening the spectral window used for the analysis (see Methods), thus decreasing the relative weight of narrow spectral perturbations compared to the total integrated spectral signal. This is illustrated in Figs. 3a-c for PEG_400_–H_2_O mixtures, where the local peak shifts from 2870 cm^−1^ to 2920 cm^−1^ as the PEG_400_ volume fraction changes from 100% to 74%. (Fig. 3a). Despite this shift, the corresponding phasor coordinates (Figs. 3b,c) remain essentially unchanged because the integration window spans a spectral range that is two orders of magnitude larger than the local shift. This demonstrates the robustness of the method with respect to moderate spectral distortions within the analyzed band.

To place the performance of spectral phasor analysis in context, we compared it with representative fitting- and regression-based approaches commonly used for Raman spectral deconvolution. We selected MCR-ALS^47-49^, a matrix-based optimization process where the fitting parameters are vectors of Raman intensities that emulate the pure spectral signatures of the constituent compounds. As representative regression approaches, we considered PCR and PLSR, which establish quantitative relationships between spectral features and sample composition through dimensionality reduction and calibration against known reference samples^50-53^. These methods differ substantially in their mathematical implementation, but all aim to infer the composition of spectrally mixed samples from Raman data.

Figures 3d–f compare the phasor framework with MCR-ALS, PCR, and PLSR, using identical experimental datasets and equivalent levels of prior information. Under these conditions, where only the fractions of the pure constituents (water and polymer) are known, spectral phasor analysis outperforms both PCR (Fig. 3e) and PLSR (Fig. 3f). For the MCR-ALS analysis (Fig. 3d), the spectrum of the pure polymer was intentionally withheld from the fitting procedure to emulate realistic biological applications, where the spectral signatures of pure components are generally unknown and the model is assumed to provide a blind, yet satisfactory approximation^47-49^. Overall, phasor analysis provides compositional estimates comparable to or more accurate than conventional approaches while avoiding iterative fitting procedures, model selection steps, and extensive calibration steps.

Next, we extend the Raman spectral phasor approach to phase-separated systems using the aqueous two-phase system (ATPS) formed by PEG and dextran. Figure 3g schematizes a phase diagram of this mixture inspired by ref. ^62^, highlighting the binodal and associated tie lines. Measurements corresponding to points (1–5) in Fig. 3h,i were performed by isolating the PEG-rich and dextran-rich phases of the ATPS. For each tie line, the polymer concentration in the coexisting phases can be independently retrieved from the initial mixture composition and the intersection of the tie line with the binodal.

Figures 3h and 3i compare the polymer volume fraction obtained from spectral phasor analysis for the dextran-rich and PEG-rich phases, respectively, with values independently measured by optical refractometry^62^. This strong quantitative agreement confirms that the phasor approach reliably reports the fractional composition of phase-separated systems. This validation establishes a benchmark that enables extension of our Raman spectral phasor analysis to single-droplet measurements.

### 2.4 Single-droplet Raman phasor analysis reveals condensate composition and small-molecule partitioning

We next applied spectral phasor analysis at the single-droplet level to characterize the composition of biomolecular condensates. As an initial model system, we examined condensates formed by oppositely charged polypeptides R_10_ and D_10_ (Fig. 4a). Focusing first on a low-frequency spectral region (1290-1470 cm^-1^) where R_10_ and D_10_ exhibit distinct Raman signatures and where water contributions are negligible, we applied the phasor approach to disentangle the respective polypeptide spectral contributions (Fig. 4b,c). This analysis revealed an asymmetric composition of the dense phase, consisting of approximately 41% positively charged R_10_ and 59% negatively charged D_10_, consistent with the intrinsically negative zeta potential of R_10_-D_10_ condensates^64^.

**Figure 4:**
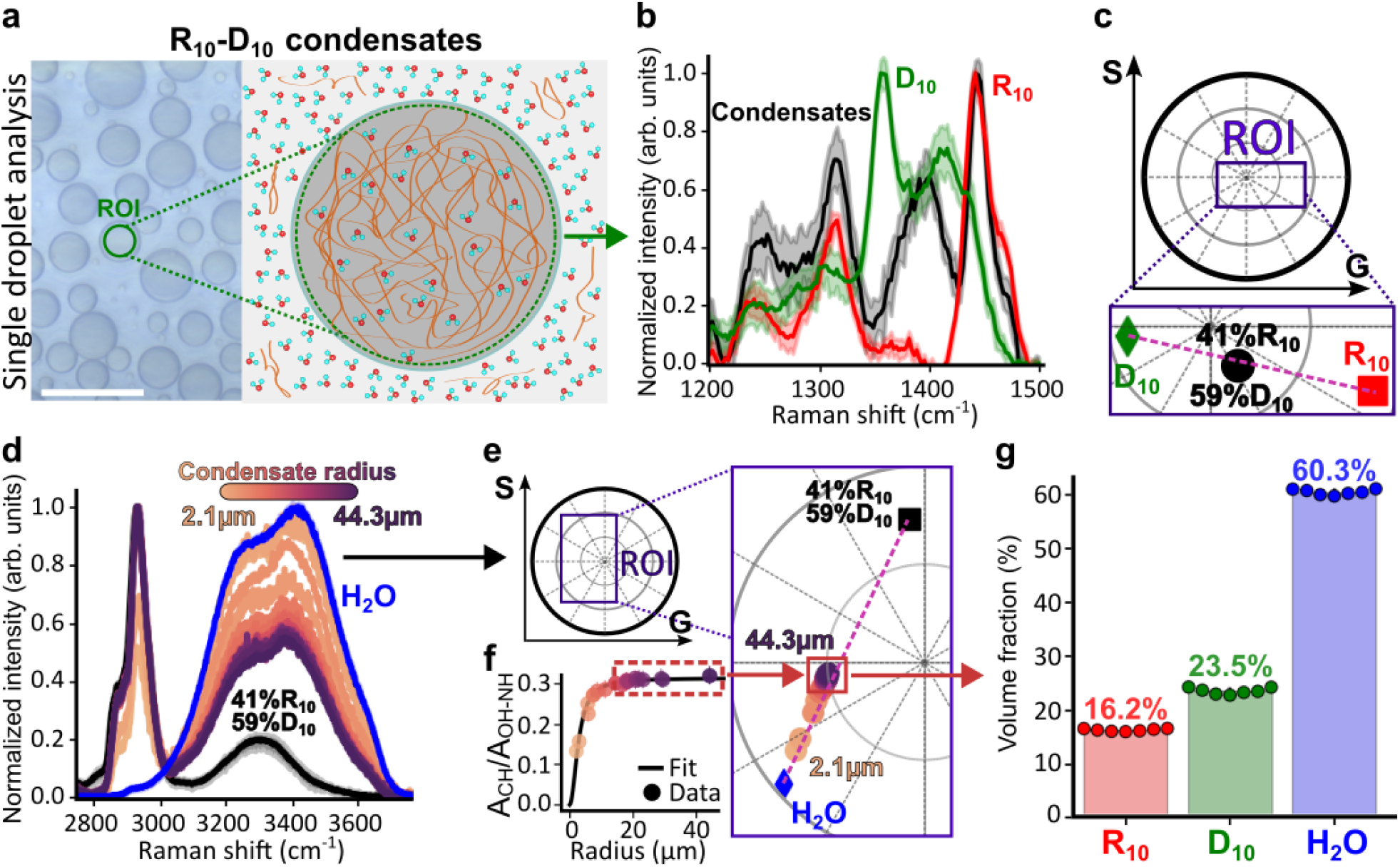
Single-droplet Raman spectral phasor analysis resolves condensate composition. **(a)** Left: bright-field image of R10-D10 condensates formed at final concentrations of 2.5 mM R10 and 2.5 mM D10. The green region of interest (ROI) indicates the Raman acquisition area. Scale bar: 40 μm. Right: schematic of protein-rich and protein-depleted phases. **(b)** Normalized Raman spectra of pure R10 (red, n = 4), pure D10 (green, n = 4) and R10-D10 condensates (black, n = 17) over the 1250-1500 cm^-1^ region. Shaded regions indicate measurement uncertainty. **(c)** Phasor representation of the spectra in (b) enabling quantification of R10 and D10 fractions using Eq. (M8). **(d)** Normalized Raman spectra of condensates of varying radii (shown in the color bar) in the 2700 cm^-1^-3800cm^-1^ region. The blue curve corresponds to pure water; the black curve is a weighed sum of the spectra of pure R10 and pure D10. **(e)** Phasor representation of the spectra in (d). The dashed purple line connects the reference points of water and polypeptide mixture. The color of each data point corresponds to that of its Raman spectrum in (d). **(f)** Ratio of integrated Raman intensities associated with C–H (2700-3100 cm^-1^) and O–H/N–H (3100-3800 cm^-1^) vibrations as a function of condensate radius *R*. The function used to fit the data was of the type *y = aR*^2^/(*R*^2^ + *b*), where *a* and *b* are fitting constants. The dashed rectangle indicates the size-independent regime. **(g)** Volume fractions of water and polypeptides in R10-D10 condensates. The data was truncated according to a threshold that corresponds to the asymptote of the A_CH_/A_OH-NH_ curve presented in (f).

We then analyzed Raman spectra acquired over the full 2800–3800 cm^−1^ region for condensates of different radii (Fig. 4d). For condensates of radii below ∼10 μm, the relative intensity of the O– H stretching band increased markedly with decreasing droplet size, indicating an increased contribution from bulk water. This effect arises because small condensates do not fully occupy the vertical extent (∼5 μm) of the confocal acquisition volume, leading to partial sampling of the surrounding dilute phase (see Fig. S4). As a result, the phasor coordinates of smaller condensates shift closer to the bulk water reference point (Fig. 4e).

Above a condensate radius of ∼10-15 μm, this size dependence vanishes as the acquisition volume becomes fully occupied by condensate material (see Fig. S4). In this regime, the ratio of integrated Raman intensities associated with C–H and O–H/N–H vibrations *A*_*CH*_/*A*_*OH*−*NH*_ reaches a plateau (Fig. 4f), defining a size-independent readout of the intrinsic protein-to-water composition of the dense phase. We therefore used this radius threshold as a cutoff for quantitative compositional analysis, yielding the volume fractions shown in Fig. 4g. Comparable analyses for all additional phase separating systems examined in this study are presented in Figs. S5-S9. Notably, because this approach resolves the polymer-to-water ratio in both the dense and dilute phases, it can in principle be employed to experimentally determine tie lines in arbitrary phase-separating system.

Finally, we investigate how small-molecule partitioning modulates condensate composition by examining the influence of ATP, a small molecule known for its physiologically multifunctional intracellular role^65, 66^. Figure 5a shows the Raman spectra of pure ATP, pure R_10_, and pure D_10_, and R_10_-D_10_ condensates formed in the presence of increasing ATP concentrations. ATP exhibits distinct spectral features within the analyzed spectral window, allowing its contribution to be resolved from those of the constituent polypeptides. As the ATP concentration increases, the condensate spectra progressively shift toward the ATP reference spectrum. Consistent with this behavior, the corresponding phasor coordinates deviate from the line connecting the pure R_10_ and D_10_ reference points and move toward the ATP phasor position (Fig. 5b). According to the geometric rules of multicomponent phasor analysis (Fig. S1), this displacement enables simultaneous quantification of the relative contributions of ATP, R_10_, and D_10_ within the dense phase.

**Figure 5:**
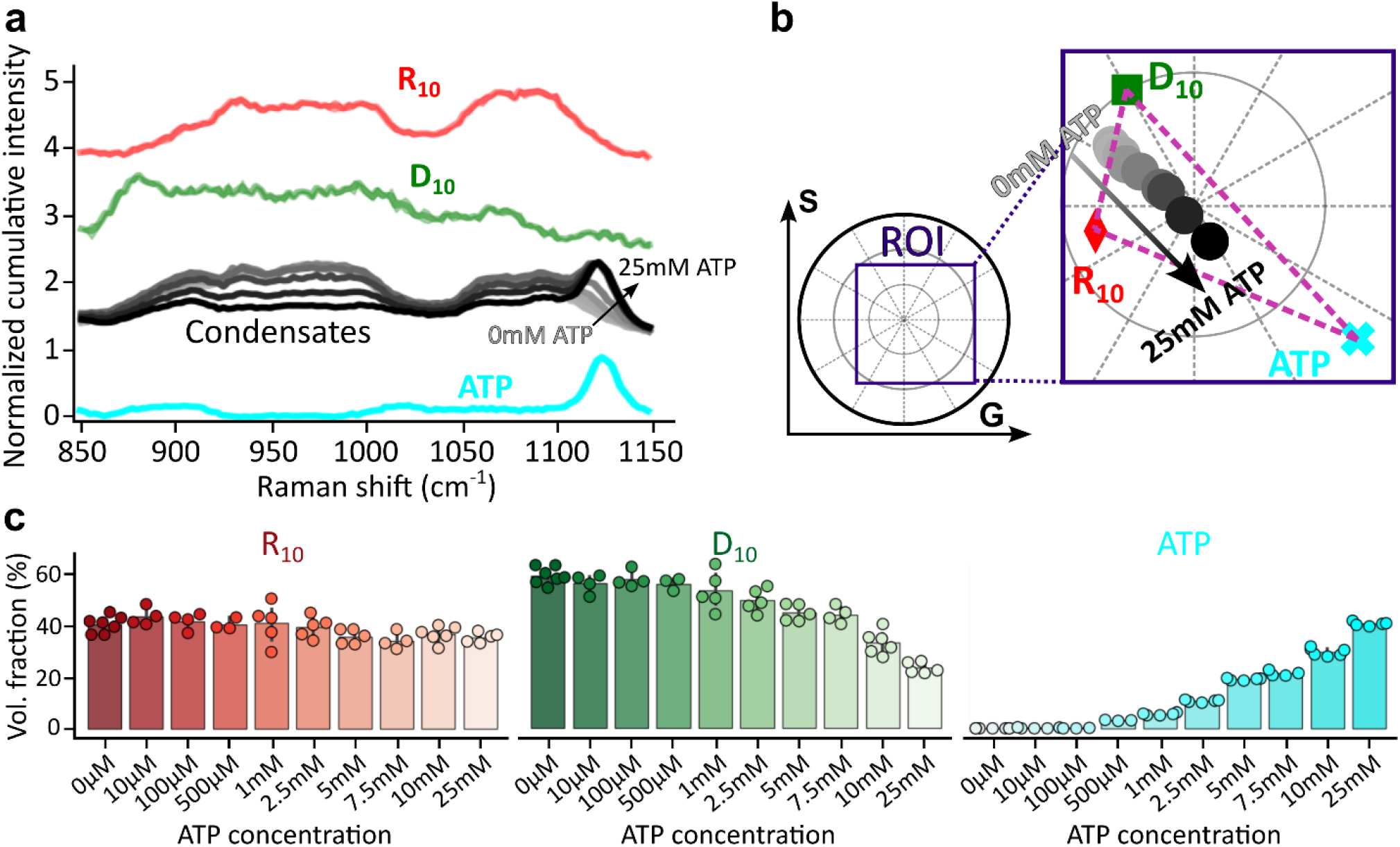
Single-droplet Raman spectral phasor analysis resolves small-molecule partitioning. **(a)** Normalized Raman spectra of pure R10 (red, n = 5), pure D10 (green, n = 5), R10-D10 condensates formed in the presence of increasing ATP concentrations (grey-black, sample numbers for each ATP concentration are given in panel c), and pure ATP (cyan, n = 5) with characteristic phosphate backbone signature at 1130 cm^−167^. **(b)** Phasor representation of the Raman spectra of pure R10 (red), pure D10 (green), pure ATP (cyan) and R10-D10 condensates (grey-black) with increasing addition of ATP (black) from 0 to 25mM. The dashed purple lines connect the reference compounds. **(c)** Phasor-based quantification of the non-aqueous molecular composition of the condensates in (b) at different ATP concentrations: 0 mM (n = 7), 0.01 mM (n = 4), 0.1 mM (n = 4), 0.5 mM (n = 3), 1 mM (n = 5), 2.5 mM (n = 5), 5 mM (n = 5), 7.5 mM (n = 4), 10 mM (n = 6) and 25 mM (n = 5).

As shown in Fig. 5c, ATP partitioning into the condensates increases monotonically with ATP concentration and reaches molecular fractions approaching 40% at 25 mM ATP. In contrast, the R_10_ fraction varies only modestly (<5%), whereas the D_10_ fraction decreases by approximately 35 percentage points, indicating preferential replacement of the negatively charged D_10_ polypeptide by ATP within the condensate phase. These results demonstrate that spectral phasor analysis enables a quantitative and label-free assessment of small-molecule partitioning and compositional remodeling within individual biomolecular condensates.

### 2.5 Raman hyperspectral phasor imaging provides spatially-resolved compositional mapping of biomolecular condensates

Figure 6 illustrates how the quantitative Raman-phasor framework introduced above can be extended to hyperspectral imaging for spatially resolved mapping of the fractional composition of biomolecular condensates at pixel resolution.

**Figure 6:**
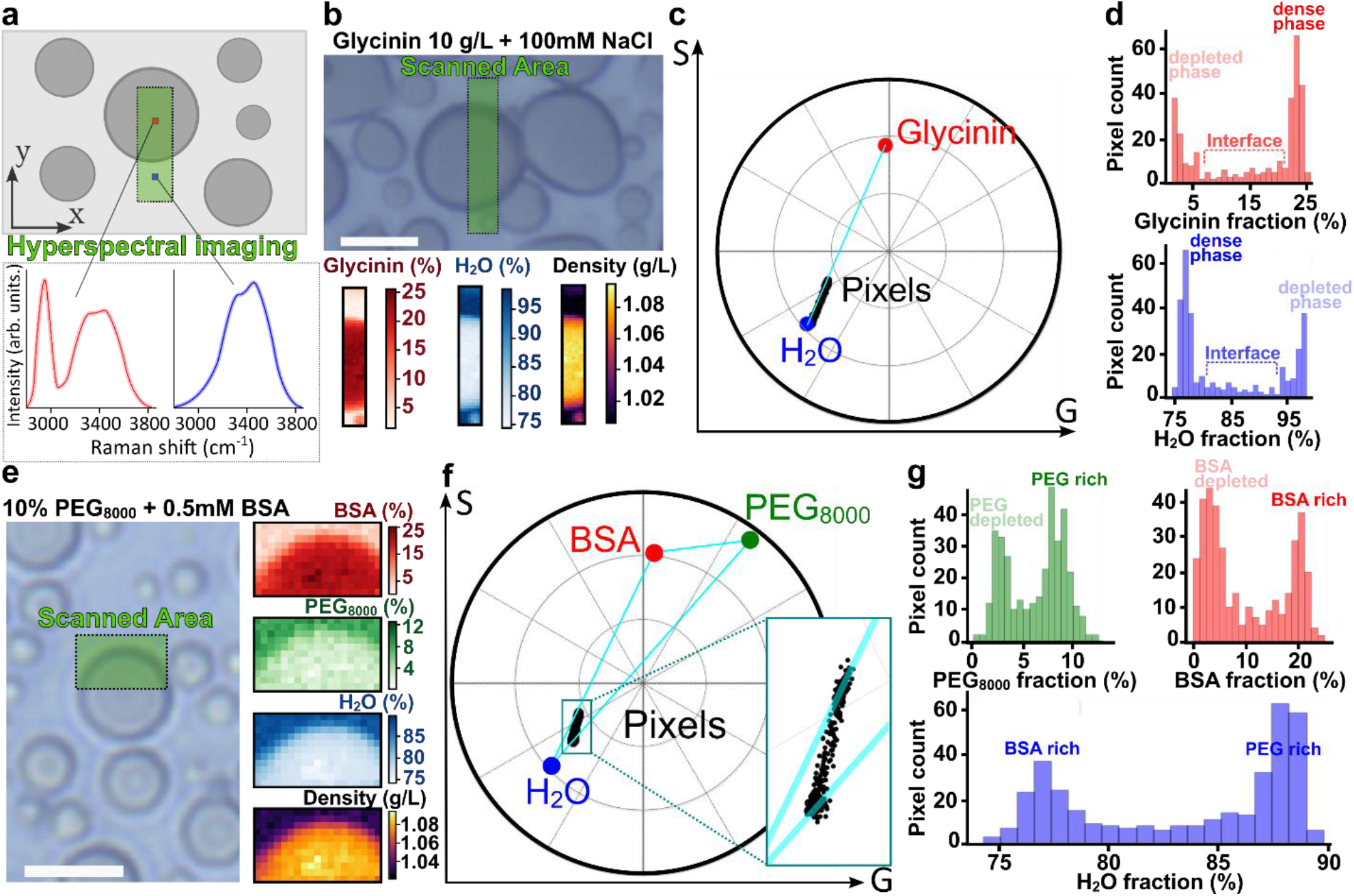
Raman hyperspectral phasor imaging enables pixel-resolved compositional mapping of biomolecular condensates. **(a)** Schematic Raman-based 2D confocal hyperspectral imaging of a condensate sample, illustrating protein-rich (dark-gray) and protein-depleted (light-gray) regions and their expected Raman spectra. **(b)** Top: bright-field image of glycinin condensates at final concentrations of 10 g/L glycinin and 100 mM NaCl (ROI outlined in green). Scale bar: 10μm. Bottom: pixel-resolved maps of glycinin (red) and water (blue) volume fractions in the ROI, and the corresponding density map (right) derived from the specific volumes of the components using Eq. (M8). Phasor representation of all pixel spectra from the ROI in (b) acquired over the 2700-3800cm^-1^ region. Pixel spectra (black dots) align along the line connecting pure glycinin (red) and pure water (blue), indicating binary mixing. (**d**) Histograms of pixel-level volume fraction of glycinin (top) and water (bottom) for the scanned area in (b). **(e)** Left: bright-field image of PEG8000-BSA condensates obtained by mixing solutions of PEG 8000 kDa and BSA buffered in PBS to the indicated final concentrations. Scale bar: 10μm. Right: pixel-resolved volume fractions maps of BSA (red) PEG8000 (green) and water (blue), and a density map computed using Eqs. (M8-M10) for the ROI indicated on the left. Phasor representation of all pixel spectra from the ROI in (e) acquired over the 2700-3800cm^-1^ region, showing ternary mixing within the triangle defined by pure BSA (red dot), pure PEG8000 (green) and pure water (blue). Histograms of the pixel-level volume fractions of PEG8000, BSA and water for the scanned area in (e).

Figure 6a,b shows that Raman hyperspectral imaging (i.e. where a full spectrum is acquired at each pixel) can be used to map the local volume fractions of glycinin protein and water across both the dense phase (glycinin condensates) and the dilute phase, based on the same phasor-based approach established in Figs. 3-5. Based on the inferred protein and water volume fractions and their specific volumes (glycinin ∼ 0.73 L/kg, water ∼ 1 L/kg, see Table S1), the local mass density can additionally be mapped down to pixel resolution (Fig. 6b, bottom). This derivation assumes that, at the concentrations explored here, the mixture density obeys the conservation relation 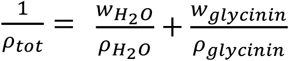 where *ρ*_*tot*_ is the pixel-resolved total density and *ρ*_*i*_ and *w*_*i*_ respectively correspond to the density and weight fraction of component *i*. In Fig. 6c, the phasor representation of spectra from all pixels in Fig. 6b shows a clear alignment along the straight line connecting pure water and pure glycinin. This confirms that within the probed spectral window, the system behaves as a strictly binary mixture of water and glycinin with spatially varying proportions.

Interestingly, the alignment observed in Fig. 6c indicates that the spectral properties of water within the dense phase are indistinguishable from those of bulk liquid water. Indeed, in a ternary scenario – involving glycinin, bulk liquid water, and a spectrally distinct population of long-lived hydrogen-bonded (“solid-like”) water molecules (as illustrated in Fig. S3 for aqueous protein solutions) – one would expect some of the data points to deviate from the water-glycinin line on the phasor plot (see Fig. S1b,c). Given the distinct Raman signatures of liquid and “solid-like” water (Fig. S3), the absence of such deviations in Fig. 6c implies that contributions from long-lived hydrogen-bonded water in the dense phase are negligible.

Figure 6d shows pixel-resolved statistical distributions of the water and glycinin volume fractions, from which dense (∼23% glycinin, ∼77% water) and dilute (∼2% glycinin, ∼98% water) phases can be clearly distinguished. Because Raman spectra are collected within a volume of constant vertical extent (i.e. volume of the focused laser beam), residual out-of-focus contributions from the surrounding dilute phase may slightly overestimate the water fraction within the dense phase, particularly near droplet boundaries (see Fig. S4). This systematic effect does not affect the alignment of the pixel-resolved spectral phasor data along the water-glycinin axis (Fig. 6c), nor the conclusion that the spectral signature of the dense phase is consistent with a binary mixture of protein and bulk-like water.

Figure 6e extends this Raman-based compositional mapping to a ternary PEG-BSA-water condensate system. The resulting maps reveal preferential enrichment of BSA within the dense phase, whereas PEG accumulates in the surrounding, more hydrated phase, consistent with previous reports on phase-separated systems of BSA and PEG of different molecular weight^46^. The corresponding phasor plot of all pixel spectra (Fig. 6f) lies within the triangle defined by the pure PEG, BSA and water references, as expected for ternary mixing. Figure 6g reports the pixel-level volume fraction distributions for PEG, BSA and water, which further resolve dense and dilute phases with PEG peaking at ∼2.5% (dense) and ∼8% (dilute), BSA at ∼20% (dense) and ∼4% (dilute), and water at ∼77.5% (dense) and ∼88% (dilute).

### 2.6 Combining Raman spectral phasors with environment-sensitive dyes reveals the compositional determinants of condensate hydrophobicity

We next relate the water-to-protein volume fractions obtained from Raman spectral analysis to the hydrophobicity of biomolecular condensates. To quantify hydrophobicity, we used the environment-sensitive dye ACDAN, whose fluorescence emission depends on the dipolar relaxation of water molecules in the immediate surrounding. By calibrating ACDAN spectral shifts against a panel of solvents with known dielectric constants, the local permittivity of a condensate can be determined at the pixel level, providing a direct measure of the condensate’s hydrophobic environment^54^. This approach enables spatially resolved mapping of water–protein interactions and allows us to correlate condensate composition, as determined from Raman spectral phasors, with their effective dielectric properties.

Figure 7a presents a Perrin-Jablonski diagram illustrating the conventional understanding^17, 19, 20^ of how ACDAN fluorescence depends on dipolar relaxation in its immediate environment: the emission redshift occurs as surrounding water molecules relax around the dye dipole. In this model, water molecules in the dense phase interact with the proteins forming the condensate, which has been proposed to slow^17, 19, 20^ dipolar relaxation through dipole-dipole interactions or suppress^44, 68, 69^ it via hydrogen bonding, or both ^70, 71^. As shown below, our results challenge the dominant role of hydrogen bonding in this process.

**Figure 7:**
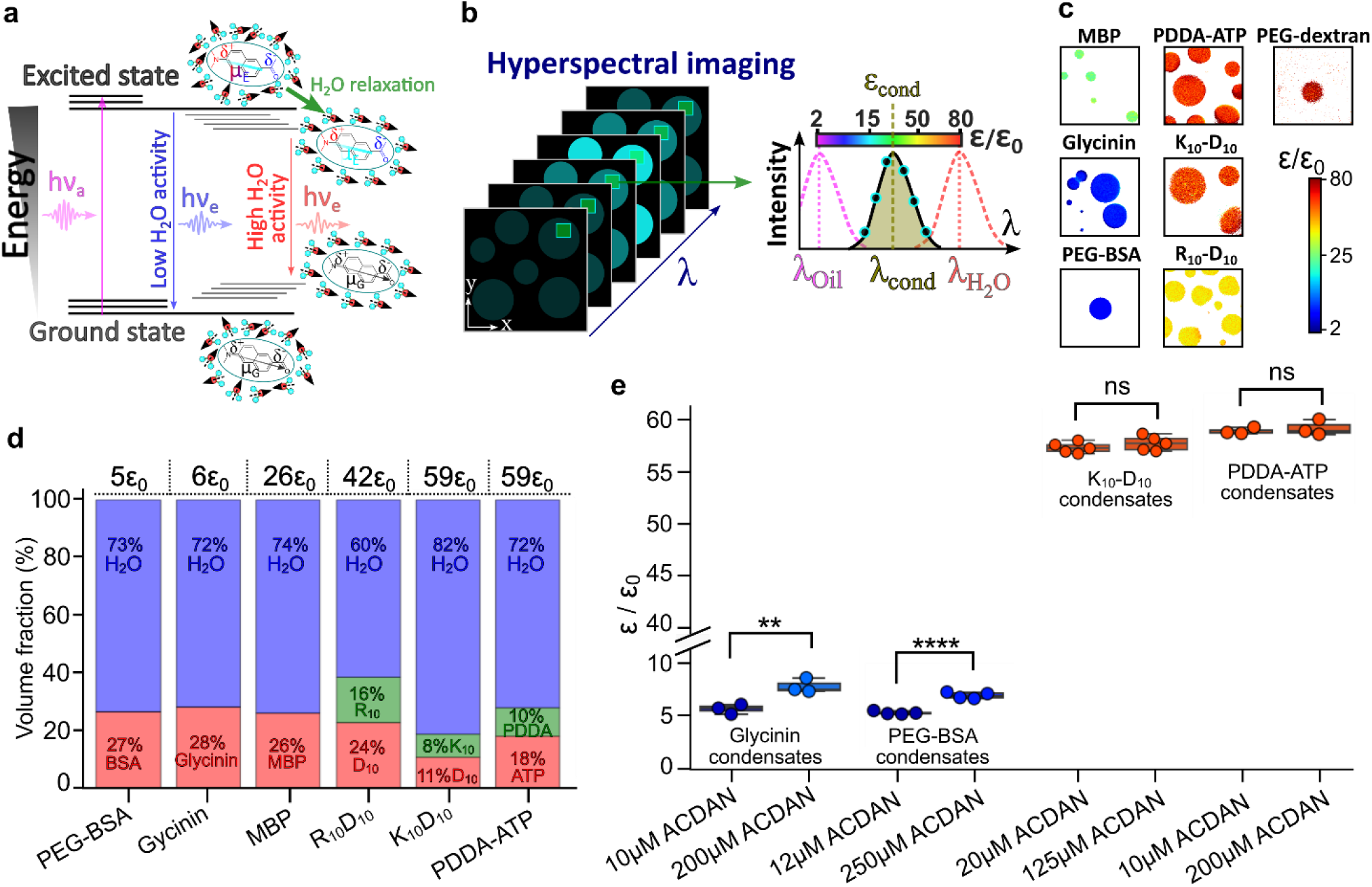
Fluorescence-based measurements of water dipolar relaxation unveil a multi-scale picture of the hydrophobicity of biomolecular condensates. **(a)** Perrin-Jablonski diagram illustrating how ACDAN fluorescence depends on dipolar relaxation of water molecules in the immediate surrounding. The molecular structure of ACDAN and its dipole moment change are sketched. Non-relaxed water dipoles cause a weak redshift, while relaxed water dipoles produce a strong redshift in the dye fluorescence. **(b)** Schematic of 2D confocal fluorescence hyperspectral imaging of a condensate sample. The local permittivity of each pixel (in the dense or dilute phase) is determined using ACDAN fluorescence calibrated against solvents of known dielectric constant^54^. **(c)** Mapping the dielectric constant of different condensate systems. The color bar shows the effective dielectric constant normalized by vacuum permittivity *ε*_0_. **(d)** Volume fraction of proteins, polymers and water in dense phases of BSA (n=8), glycinin (n=6), MBP (n=3), R10-D10 (n=7), K10-D10 (n=6) and PDDA-ATP (n=8) condensates. The effective dielectric constants indicated atop each bar were obtained from ACDAN hyperspectral imaging in (c). **(e)** Comparison of measured dielectric constants at low and high ACDAN concentrations. Differences at high dye concentration reflect partial saturation of protein surfaces and exposure of the dye to water, not changes in water partitioning in the condensates. The number of stars correspond to the negative power of 10 associated with the p-value of the Student’s t-test.

Using confocal hyperspectral imaging calibrated with ACDAN^54^, we determined the effective dielectric properties of condensates as illustrated in Fig. 7b, revealing a broad continuum of hydrophobicity across condensates formed by LLPS of various proteins and polymers (Fig. 7c). While ACDAN was chosen for its robust and broadband dielectric measurement characteristics^54^, Fig. S10 shows that these trends are well reproduced using Nile Red as an independent environment-sensitive fluorescent reporter^72, 73^, demonstrating the generality of this classification.

Figure 7d presents the compounds volume fractions for condensates ordered from high to low hydrophobicity. Notably, in short and unstructured oppositely charged polypeptide condensates, e.g. R_10_-D_10_ versus K_10_-D_10_, the water volume fraction, respectively 60% and 82%, strongly correlates with the dielectric constant, 42*ε*_0_ and 59*ε*_0_ measured with ACDAN. Interestingly, condensates composed of larger proteins (glycinin, BSA, MBP) exhibit high water content (> 72%) while showing extremely weak dipolar relaxation (high apparent hydrophobicity), while PDDA-ATP condensates are highly hydrophilic despite similar water content, consistent with values reported in the literature^28, 74^. Additional results presented in Fig. S11 at different PDDA concentrations confirm that these effects are not dominated by the PDDA molecule itself.

The above observations indicate that water partitioning alone does not fully account for the range of measured dielectric constants. Importantly, Figs. 6, S3, S5-S9 show that the fraction of hydrogen-bonded water in all dense phases is negligible, ruling out suppressed dipolar relaxation via hydrogen bonding. Instead, these trends are consistent with a slowdown of water dynamics induced by protein hydration, as previously reported by time resolved infrared spectroscopy^75^, terahertz spectroscopy^58, 69^, and dielectric relaxation spectroscopy^70^, which limits dye dipolar relaxation and thus emission redshift^76^.

We next considered whether the dielectric properties measured in condensates formed by large globular proteins could be influenced by an uneven spatial distribution of the ACDAN probe. In homogeneous polymer solutions such as aqueous PEG, ACDAN-reported dielectric constants closely follow predictions from effective-medium theory^77^, consistent with uniform probe mixing (Fig. S12a). In contrast, aqueous solutions of glycinin and BSA show systematic deviations from the Maxwell–Garnett model^77^ when protein volume fractions determined by Raman phasor analysis are used as input (Fig. S12b,c). These deviations indicate that ACDAN does not sample the mixture uniformly, but preferentially partitions into structurally heterogeneous, protein-rich regions.

In condensates formed by these proteins, such uneven mixing provides a complementary explanation for the strong suppression of dipolar relaxation inferred from ACDAN fluorescence. As shown in Fig. 7e, increasing the ACDAN concentration leads to a somewhat higher apparent dielectric constant, consistent with partial saturation of water-depleted structural regions and increased exposure of the dye to the surrounding aqueous environment (note that this effect is absent in hydrophilic, unstructured systems such as PDDA-ATP condensates). Importantly, Raman spectroscopy demonstrates that this effect is not accompanied by enhanced water partitioning into the dense phase. Specifically, the ratio of C–H to O–H/N–H vibrational intensities *A*_*CH*_/*A*_*OH*−*NH*_ remains systematically higher at elevated ACDAN concentrations (Fig. S13), indicating that the protein-rich phase does not become more hydrated. Together, these results show that the dielectric response reported by ACDAN in structured protein condensates reflects both slowed water dynamics near hydrated protein backbones and probe sensitivity to hydrophobic pockets, rather than changes in bulk water content. This interpretation is consistent with numerical studies of water dynamics reporting orders-of-magnitude increases in water relaxation times near structured protein surfaces^21-23^.

## 3. DISCUSSION

The present study provides a direct *in situ* means of quantifying the composition of biomolecular condensates, bypassing invasive procedures such as centrifugation and drying that are commonly used to estimate condensate hydration levels^28, 29^ and client molecule partitioning^30^. These conventional approaches involve substantial variations in temperature and pressure, which are likely to perturb the native phase behavior of condensates. In contrast, the Raman-based framework introduced here enables label-free, quantitative measurements at the single-droplet level, allowing precise determination of protein and water volume fractions without the need for fluorescent tagging. This is particularly important given that molecular labels can alter protein topology, intermolecular interactions, and ultimately partitioning efficiencies and binding^78-80^. This study establishes Raman spectral phasor analysis as a powerful and minimally perturbative tool for resolving protein composition with high precision at the single-condensate level.

In the current literature, Raman-based characterization of condensate composition has relied heavily on fitting and regression methodologies^36, 38, 39, 45, 49^, particularly for in vivo systems, where the molecular composition is intrinsically heterogeneous and incompletely known. In this context, the spectral phasor approach provides a computationally lightweight strategy for resolving the molecular composition of biomolecular condensates based on the linear algebraic properties of mixtures in Fourier space.

In the context of in vivo condensates, this framework could be used to distinguish between broad classes of biomolecular constituents, such as proteins, nucleic acids, lipid-rich domains, and cytosolic components, provided that representative spectral signatures are available as reference points in phasor space. Importantly, the objective need not be the exhaustive identification of every molecular species present. Rather, the approach enables the quantification of the relative contribution of selected molecular classes or target molecules within an otherwise complex intracellular environment.

Alternatively, the spectral phasor formalism can be used to quantify the partitioning of a specific molecular species within heterogeneous or evolving environments by comparing native and perturbed states of the system. In this representation, compositional changes correspond to trajectories in phasor space, whose projection onto a reference axis follows a lever-rule-like behavior (Fig. S1), enabling direct estimation of partition coefficients without explicit spectral fitting.

Finally, extending the present framework to multi-harmonic spectral phasor analysis (*n* > 1; see Methods) may provide additional spectral degrees of freedom that improve discrimination between chemically similar species^81^. Combined with emerging Raman reference libraries and advances in intracellular Raman imaging, this could offer a route toward increasingly resolved compositional mapping of biomolecular condensates in living cells.

Beyond its methodological advantage, this study provides a new perspective on a long-standing biophysical question about the role of water dynamics in shaping the macroscopic properties of crowded macromolecular environments, and in particular regarding the contribution of hydrogen bonding in defining condensate hydrophobicity. Previous Raman studies have reported that the water structure and hydrogen bonding depend on the type of LLPS (i.e. segregative versus associative) and on water interactions with the protein scaffold^44, 57^. Our results contrast these interpretations, and we identify two key factors that reconcile this discrepancy. First, earlier analysis did not account for the influence of condensate size, which we show to significantly influence the measured Raman signal (Figs. 4, S5-S10). Second, the intrinsic spectral contributions of O–H and N–H bonds in the protein backbone were not explicitly considered^44, 57^, potentially leading to inadequate attribution of spectral changes in the O–H vibration region to water alone (Fig. 2c). Once these contributions are properly accounted for, the apparent deviations in the Raman spectral profile of condensate water disappear, revealing a spectral signature that is indistinguishable from bulk liquid water even in condensates that appear highly hydrophobic when probed by fluorescence-based methods (Fig. 7). This observation is consistent with previous measurements indicating that only a minor fraction (∼10%) of intracellular water exhibits significantly slowed dynamics in highly crowded environments^75^ and suggests that bulk-like water remains the dominant solvent population within condensates.

Importantly, our combined Raman and fluorescence measurements reveal that neither water content nor specific chemical features of the macromolecule scaffold alone can account for the wide range of effective dielectric constants observed across different condensate systems (Fig. 7). Instead, condensate hydrophobicity probed via fluorescent reporters emerges from a more complex interplay between water partitioning, protein structure, and local hydration dynamics. While water fraction strongly influences this dielectric response in condensates formed by short, unstructured polypeptides, condensates composed of larger, structured proteins exhibit extreme apparent hydrophobicity despite retaining a high water content (>72%). Moreover, our Raman analysis shows that long-lived hydrogen-bonded water molecules contribute negligibly to the composition of the dense phase across all systems studied, ruling out suppressed dipolar relaxation via hydrogen bonding as the dominant mechanism. These findings are consistent with two-dimensional infrared spectroscopy reports of low hydrogen-bond lifetimes between water and protein molecules inside condensates (below picoseconds^82^), and an overall slowdown of water dynamics near protein surfaces, as previously observed by time resolved infrared spectroscopy, terahertz and dielectric relaxation spectroscopy^58, 69, 70, 75^, which can strongly affect the emission response of environment-sensitive dyes^76^.

Finally, these results suggest a conceptual picture in which condensates can act as heterogeneous protein hubs where environments of different hydrophobicity coexist at different length scales. Structured regions within protein-rich phases may locally enrich highly hydrophobic small molecules, while more hydrophilic species preferentially reside within the remaining water-rich fraction of the condensate.

More broadly, the present framework establishes a route toward non-invasive characterization of condensate microarchitecture by linking local dielectric measurements to quantitative compositional mapping. An important next step will be to determine whether the apparent decoupling between condensate composition and fluorescence-reported hydrophobicity originates from structurally distinct microenvironments within the dense phase. Perturbations that alter scaffold organization, such as changes in protein structure, intermolecular crosslinking, or also the introduction of amino acids and small molecules known to modulate intermolecular interactions^83-85^, are expected to remodel these microenvironments and thereby alter both client partitioning and local dielectric responses. Combining environment-sensitive fluorescence reporters with quantitative Raman-phasor compositional analysis should provide a promising strategy for directly linking condensate composition, microarchitecture, and client chemistry.

Such multiscale hydrophobic organization has other important implications for condensate function, particularly at interfaces. Differences in dipolar relaxation between condensates and their surrounding aqueous environment are expected to influence interfacial tension and wetting behavior, thereby shaping the assembly of core-shell multifunctional compartments^4, 8, 86^. Recent studies further indicate that these hydrophobicity contrasts play an important role in condensate– membrane interactions and mutual remodeling^8, 35, 54^, processes central to cellular functions such as endocytosis^87^ and membrane repair^88^. More generally, the combined approach of Raman microscopy and fluorescence imaging with an environment-sensitive dye presented here enables robust compositional mapping of condensates formed by proteins with diverse sizes, structures, and phase-separation mechanisms, providing a quantitative framework for investigating the physical chemistry of biomolecular condensates in both biological and bioengineered systems.

## 4. METHODS

### 4.1 Materials

6-acetyl-2-dimethylaminonaphthalene (ACDAN) was purchased from Santa Cruz Biotechnology (USA). Polydiallyldimethylammonium chloride (PDDA, 200-350 kDa, 20 wt% solution in water), adenosine triphosphate (ATP) and sodium hydroxide (NaOH) were obtained from Sigma-Aldrich (Missouri, USA). Poly(ethylene glycol) (PEG 8000, Mw 8 kg/mol lot #MKBT7461V; PEG 3350, Mw 3.350 kg/mol lot #110K0169 and PEG 400, Mw 0.4 kg/mol) and dextran from Leuconostoc mesenteroides (molecular weight between 400 kDa and 500 kDa) were purchased from Sigma-Aldrich. The oligopeptides, poly-L-lysine hydrochloride (degree of polymerization, n = 10; K10), poly-L-arginine hydrochloride (degree of polymerization, n = 10; R10) and poly-L-aspartic acid sodium salt (n = 10; D10) were purchased from Alamanda Polymers (AL, USA) and used without further purification (≥95%). BSA (≥98% purity, 66 kg/mol), MBP bovine (≥90%) and Polyvinyl alcohol (PVA, Mw 145 kg/mol) were purchased from Merck (Darmstadt, Germany). All solutions were prepared using ultrapure water from SG water purification system (Ultrapure Integra UV plus, SG Wasseraufbereitung) with a resistivity of 18.2 MΩ cm.

### 4.2 Condensate formation and labelling

Coverslips for confocal microscopy (26 × 56 mm, Waldemar Knittel Glasbearbeitungs GmbH, Germany) were washed with ethanol and water, then passivated with a 40 mg/mL PVA solution and an aliquot of 10 μL of condensates suspension was placed on the coverslip before imaging.

#### PDDA-ATP condensates

Phase separated droplets were formed by gently mixing aliquots of stock solutions of ATP and PDDA (in this order) with ACDAN or Nile Red in pure water to a final volume of 10 μL. The final concentration of each component was as follows: 14.8 mM ATP, 4.9 mM PDDA, 10 μM ACDAN and 60μM Nile Red.

#### Glycinin condensates

Freeze-dried glycinin was a gift from Dr. Nannan Chen. The purification is detailed in ref ^89^. A 20 mg/mL glycinin solution at pH 7 was freshly prepared in ultrapure water and filtered with 0.45 μm filters to remove any insoluble materials. To form the condensates, the desired volume of the glycinin solution was mixed with the same volume of a NaCl solution of twice the desired final concentration. In this manner, the final protein concentration was 10 mg/mL ^89^. In the case of fluorescence-based measurements, the final concentration of ACDAN and Nile Red were 5 μM and 60 μM respectively.

#### K_10_-D_10_ and R_10_-D_10_ condensates

Phase separated droplets were formed by gently mixing aliquots of stock solutions of D_10_ and K_10_ or R_10_ (in this order) with ACDAN or Nile Red in pure water to a final volume of 10 μL. The final concentration of each component was as follows: 2.5 mM D_10_, 2.5 mM K_10_ or R_10_, and 15 μM ACDAN and 60μM Nile Red in the case of fluorescence-based assays. In the ATP partitioning assays in R_10_-D_10_ condensates, ATP was added to the aqueous solution prior to addition of the polypeptides, such that condensates formed in the presence of ATP.

#### PEG-BSA condensates

Phase separated droplets were formed by gently mixing aliquots (via pipetting and releasing 3 times the total volume) in a 1:1 ratio of stock solutions of 20% PEG-8000 in phosphate buffer saline (PBS) to which 30 μM ACDAN or 60μM Nile Red were added for fluorescence experiments with a 1mM BSA solution dissolved in PBS to a final volume of 10 μL.

#### PEG-Dextran condensates

A polymer solution in composed of the desired weight fractions of PEG and dextran were prepared and left, when relevant, for 2 days to completely phase separate and equilibrate. In the case of fluorescence measurements ACDAN was then added to each phase to reach a final concentration of 25 μM.

#### MBP condensates

MBP condensates were prepared by following the procedure described in ^90^. Briefly, a 5 mg/mL solution of MBP dissolved in water was mixed with a 20 mM NaOH solution of 10 μM ACDAN in a 1:1 ratio.

### 4.3 Confocal microscopy and fluorescence hyperspectral imaging

Hyperspectral images were acquired using a confocal Leica SP8 FALCON microscope equipped with a 63×, 1.2 NA water immersion objective (Leica, Mannheim, Germany). The microscope was coupled to a pulsed Ti:Sapphire laser MaiTai (SpectraPhysics, USA), with a repetition rate of 80 MHz. A two-photon wavelength of 780 nm was used for ACDAN excitation, and a DPSS laser of wavelength 561nm was used for Nile Red excitation. Image acquisition was performed with a frame size of 512 × 512 pixels ^35, 91^ and a pixel size of 72 nm×72 nm using a HyD SMD detector in standard mode. For hyperspectral imaging, the xyλ configuration of Leica SP8 was used, sequentially measuring in 32 channels with a bandwidth of 9.75 nm in the range from 416 to 728 nm. Some hyperspectral images were realigned using the ImageJ software and all hyperspectral stacks were processed into .r64 files by the SimFCS software developed at the Laboratory of Fluorescence Dynamics (available at no cost at https://www.lfd.uci.edu/globals/), and analyzed using Python code based in the PhasorPy library (available at https://www.phasorpy.org/).

### 4.4 Raman confocal analysis and hyperspectral imaging

Raman images and spectra were acquired with a Raman confocal microscope Alpha300 R (WITec GmbH, Germany) with Zeiss EC Epiplan 50×/0.75 objective, at an excitation wavelength of 532 nm and 50 mW laser power. Spectra were acquired in the range 400–4100 cm^−1^. The Raman band of the silicon wafer was used to calibrate the spectrometer. Data were acquired using a 3 s exposure time with four accumulated scans per measurement, which were averaged prior to be further analysed using Project FIVE v.5.2 data evaluation software from WITec. All acquired spectra and hyperspectral scans were corrected for cosmic rays and the baseline was subtracted using a built-in background subtraction shape function of empirically optimized shape size of 500.

The hyperspectral scans were further processed using Python programming code for converting the Raman signal sampled in individual pixels into phasor space coordinates.

### 4.5 Spectral phasor analysis of Raman and fluorescence hyperspectral imaging data

Spectral phasor analysis was applied to both fluorescence and Raman hyperspectral imaging data to provide a unified, model-free representation of spectral heterogeneity at the pixel level. In this approach, each measured spectrum is mapped onto a point in a two-dimensional phasor space defined by the real (G) and imaginary (S) components of the first harmonic of its Fourier transform. For a given pixel, the phasor coordinates (G, S) are defined as:

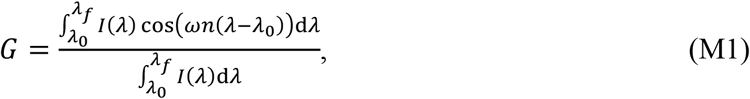

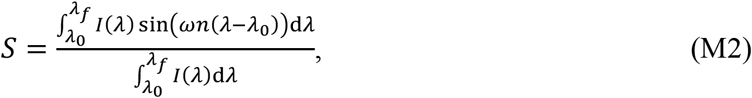

where *I*(*λ*) denotes the measured spectral intensity as a function of the detection coordinate *λ*, and (*λ*_0_; *λ*_*f*_) defines the spectral window of acquisition. This range depends on instrumental constraints and the type of detector used for the analysis. Here, *λ* is used as a generic spectral coordinate, applicable to both fluorescence emission spectra and Raman spectra. For Raman measurements, *λ* maps the Raman shift within the restricted spectral ranges analyzed, allowing the same formalism to be used.

In the present work, the spectral range 416–728 nm was used for ACDAN fluorescence data. For Raman measurements, truncated portions of vibrational spectrum spanning approximately 1050– 3850 cm^−1^ were analyzed, corresponding to the fingerprint and C–H stretching regions. Note that changing the detection range will necessarily result in a change of the relative positions of different points on the phasor plot; therefore, the same detection range was used consistently across all experiments to enable quantitative comparison. The harmonic number *n* determines which Fourier harmonic of the spectrum is used to calculate the phasor coordinates. The harmonic number n represents the expected degree of nonlinearity of the analysed signal, because we neglect non-linear effects in our approach, we set *n*=1. Equivalently, *n* is the number of cycles that the trigonometric function covers over the wavelength range, in units of the angular frequency ω:

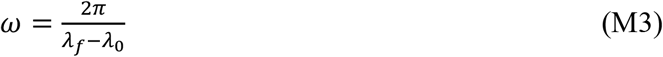

When imaging with a microscope, we acquire a finite number of spectral steps corresponding to the number of detection windows that cover the spectral range. The continuous expression in Eqs. (M1, M2) are therefore expressed as a discretized approximation:

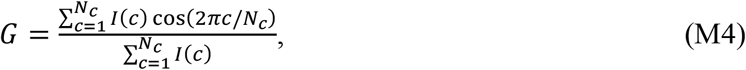

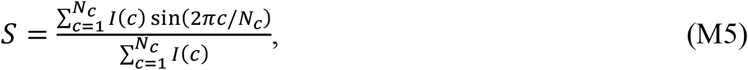

where *I*(*c*) is the pixel intensity measured in channel *c* and *N*_*c*_ is the total number of spectral channels. Although the total number of channels is finite (32 for fluorescence and over 200 for Raman data), the resulting phasor coordinates S and G can be treated as quasi-continuous due to the high photon counts per pixel and channel (>10^2^).

A key property of the spectral phasor representation is its linearity: the phasor of a spectrum composed of multiple independent components lies at the vectorial linear combination of the phasors of the individual components. Consequently, mixtures of two spectral species fall along the straight line connecting their pure-component phasors, and the relative contribution of each component can be inferred from their position along this line.

Note that the phase angle *ϕ* and the modulation *M* of each phasor are given by:

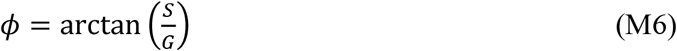

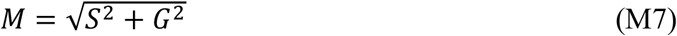

### 4.6 Geometric computation of fractional compositions based on phasor space coordinates

Using the spectral phasor coordinates of pure reference compounds (e.g. water and lyophilized polymers, see Fig. S1b), the volume fraction occupied by a given compound in a mixed spectrum can be computed geometrically. When reference mixtures of known composition are used for calibration, this fractional contribution provides a quantitative readout of the component volume fraction. For a binary mixture of components a and b, the volume fraction of component b is given by

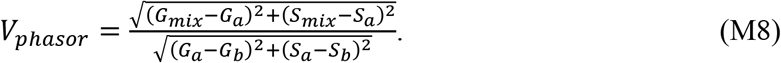

Here, (*G*_*a*_, *S*_*a*_), (*G*_*b*_, *S*_*b*_) are the phasor coordinates of the two reference components *a* and *b* (e.g. polymer and water), and (*G*_*mix*_, *S*_*mix*_) is the phasor of their mixture. By construction, *V*_*phasor*_ *=* 0 corresponds to a pure component *a*, whereas *V*_*phasor*_ *=* 1 corresponds to a pure component *b*. The fractional contribution of component *a* is therefore given by 1 − *V*_*phasor*_.

A similar equation was used in the case of 3 pure compounds (see Fig. S1c). However, in such case, the point of coordinate (*G*_*a*_, *S*_*a*_) is taken as the intersection between the straight line connecting the data point of coordinates (*G*_*D*_, *S*_*D*_) and the pure compound of interest (*G*_*b*_, *S*_*b*_) with the straight line connecting the two other compounds of coordinates (*G*_*c*_, *S*_*c*_) and (*G*_*d*_, *S*_*d*_). The expression of the coordinates *G*_*a*_ and *S*_*a*_ for a 3-compound system consequently reads:

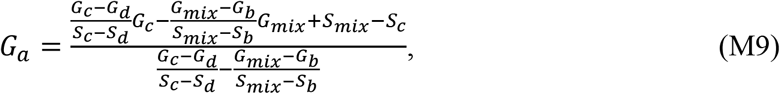

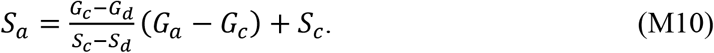

## 4 ACKNOWLEDGEMENTS

A.M. acknowledges support from Alexander von Humboldt Foundation and CONICET. We acknowledge Nannan Chen for providing glycinin. We also acknowledge support from the German Academic Exchange Service (Deutscher Akademischer Austauschdienst, DAAD) in the framework of project 57654674. R.D. acknowledges the ComeInCell network funded by the European Union’s Horizon Europe research and innovation program under the Marie Sklodowska-Curie grant agreement No. 101168939. We acknowledge Luca Bertinetti for his proofreading of the manuscript and insightful comments.

