## Supplementary Information for "Seeing the chemistry of biomolecular condensates: *in situ* mapping of composition and water content"

E. Sabri[1,*], A. Mangiarotti[1,2,3], C.N.Z. Schmitt[1] and R. Dimova[1,*]

[1]*Max Planck Institute of Colloids and Interfaces, Science Park Golm, 14476 Potsdam, Germany.*
[2]*Centro de Investigaciones en Química Biológica de Córdoba (CIQUIBIC), CONICET, X5000HUA Córdoba, Argentina.*
[3]*Departamento de Química Biológica Ranwel Caputto, Facultad de Ciencias Químicas, Universidad Nacional de Córdoba, X5000HUA Córdoba, Argentina.*

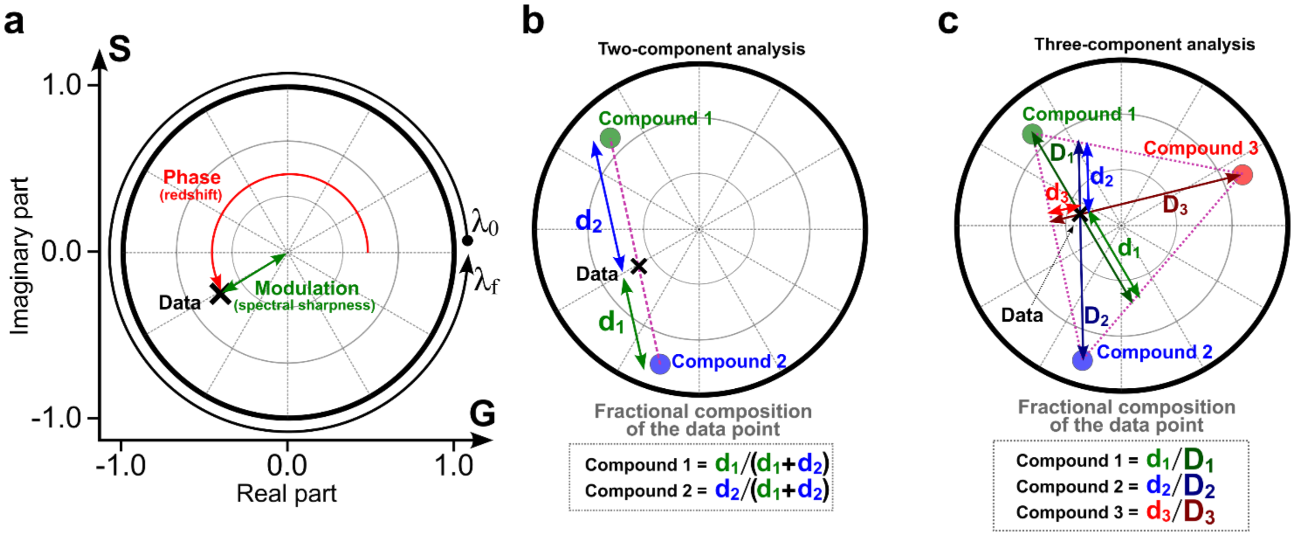


**Figure S1: Spectral deconvolution of Raman signal in phasor space allows to quantify the fractional composition of pure components. (a)** Illustration of the 2D representation of a single spectrum (black cross) on a given spectral interval $[\lambda_0; \lambda_f]$ in phasor space. The phase (red) represents the angular coordinate and encodes the effective redshift of the spectrum, and the modulation (green) represents the radial coordinate and encodes the effective sharpness of the spectral profile on the $[\lambda_0; \lambda_f]$ interval. **(b)** Illustration of a two-component deconvolution in phasor space. Assuming that the data point representing the spectrum of interest is measured from a weighed sum of the spectra of compounds 1 (green dot) and 2 (blue dot) in a priori unknown proportions, the fractional composition of the measured data can be retrieved from the geometric ratios referenced in the bottom panel. **(c)** Illustration of a three-component deconvolution in phasor space. Assuming that the data point representing the spectrum of interest is measured from a weighed sum of the spectra of compounds 1 (green dot), 2 (blue dot) and compound 3 (red dot) in a priori unknown proportions, the fractional composition of the measured data can be retrieved from the geometric ratios referenced in the bottom panel.

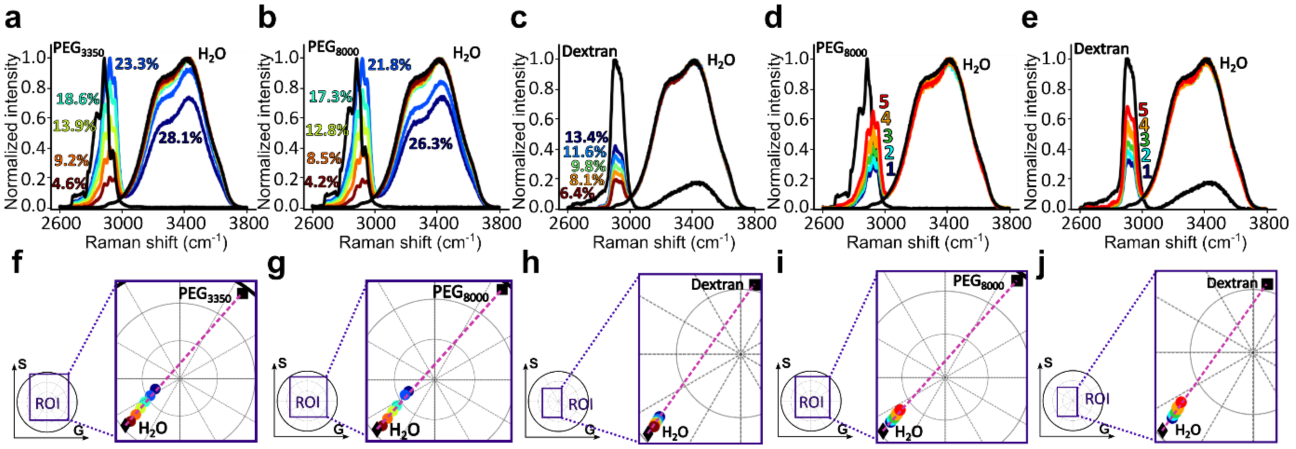


**Figure S2: Raman spectra and phasor space representations of PEG and dextran aqueous solutions. (a-e)** Raman spectra of (a) $PEG_{3350}$-water mixtures (28.1%, 23.3%, 18.6%, 13.9%, 9.2%, 4.6% $PEG_{3350}$ volume fraction) (b) $PEG_{8000}$-water mixtures (26.3%, 21.8%, 17.3%, 12.8%, 8.5%, 4.2% $PEG_{8000}$ volume fraction) (c) Dextran-water mixtures in different volume mixing ratios (13.4%, 11.6%, 9.8%, 8.1%, 6.4% dextran volume fraction). **(d-e)** PEG-rich phase and dextran-rich phase of PEG-dextran aqueous two-phase systems. Points 1-5 correspond to the phase-diagram points 1-5 measured in Fig. 3d-f. (f-j) Phasor space representation of the spectra shown respectively in (a-e), the dashed lines are guides to the eye linking the water and polymer spectral phasor coordinates.

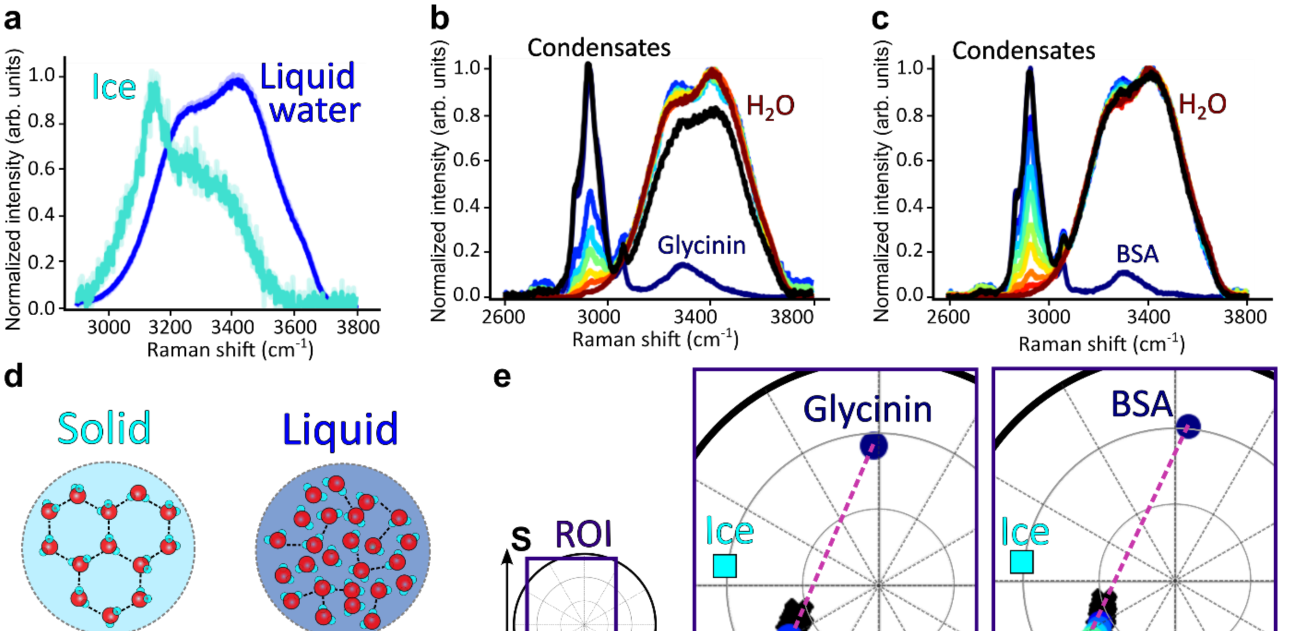


**Figure S3: Water hydrogen bonding does not play a role in the apparent hydrophobicity of biomolecular condensates. (a)** Raman spectra of ice (cyan) and liquid water (blue) over the (2900-3800$cm^{-1}$ region). **(b)** Raman spectra of glycinin-water mixtures with protein volume fractions of 100% (dark blue), 15.4% (blue), 11.4% (cyan), 7.5% (green), 3.7% (yellow), 1.5% (red) and 0% (dark red). The black curve represents the averaged spectrum of glycinin condensates (obtained by mixing a glycinin aqueous solution with a NaCl solution to 10g/L glycinin and 100mM NaCl final concentrations) for large enough condensates so that the acquisition volume (see Fig. S3) would be filled with condensate material and spectral profiles are no longer affected by condensate radius. **(c)** Raman spectra of BSA-water mixtures with protein volume fractions of 100 % (dark blue), 25.9% (blue), 23.8% (turquoise), 19.5% (cyan), 15.4% (lime green), 11.4% (green), 7.5% (yellow), 3.7% (orange), 1.8% (red), 0% (dark red). The black curve represents the averaged spectrum of BSA condensates (obtained by mixing a BSA aqueous solution with a $PEG_{8000}$ solution to 0.5 mM BSA and 10% weight fraction $PEG_{8000}$ concentrations) for large enough condensates so that the acquisition volume (see Fig. S3) would be filled with condensate material and spectral profiles are no longer affected by condensate radius. **(d)** Illustration of solid and liquid water phases at the microscopic-scale. In the solid state (left panel), water exhibits a highly stable hydrogen bonded water molecule network whereas in the liquid state (right panel) the interatomic bond population is dominated by covalent bonding. These trends can be assessed based on the redshifted peak appearing in the Raman spectrum of ice presented in (a), where the weaker nature of hydrogen bonds versus covalent bonds yields a redshifted overall spectrum. **(e)** Phasor space representation of the spectra in respectively (b-c), the dotted lines are guides to the eye linking the water and protein spectral phasor coordinates. The spectrum of ice water is represented as a square cyan datapoint and serves as a landmark hydrogen bonded water.

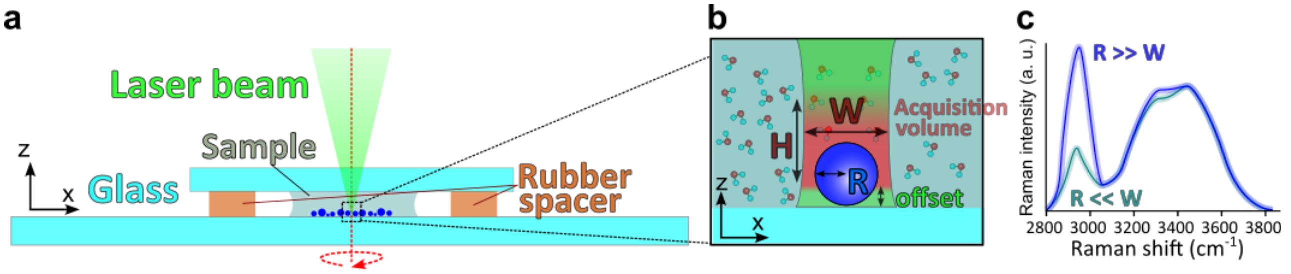


**Figure S4: Finite confocal acquisition volume introduces a condensate-radius dependence in single-droplet Raman spectra. (a)** Illustration of the experimental observation chamber. The red dotted line and arrow illustrate the optical axis, about which the system is rotationally symmetric. **(b)** Zoomed-in illustration of the imaged portion of the sample. The red, green and blue regions respectively represent the effective confocal acquisition volume, the laser beam with beam waste W, and a condensate droplet of radius R. **(c)** Schematic Raman spectra for condensates with radii much smaller (turquoise) and much higher (blue) that the characteristic dimensions of the acquisition volume. For small droplets, partial filling of the acquisition volume enhances the relative contribution of the surrounding aqueous phase.

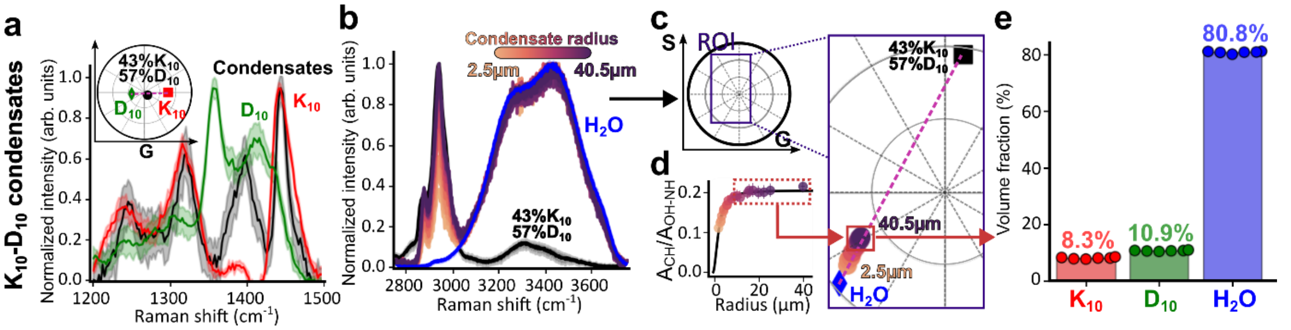


**Figure S5: Single-droplet phasor analysis of $K_{10}$-$D_{10}$ condensates. (a)** Normalized Raman intensity over the high-frequency part of the signature region. The red, green and black curves correspond to the Raman spectra of pure $K_{10}$, pure $D_{10}$ and $K_{10}$-$D_{10}$ condensates. The top left inset shows the phasor representation of the Raman spectra of pure $K_{10}$, $D_{10}$ and condensates. The proportions of $K_{10}$ and $D_{10}$ were computed based on Eq. M1. **(b)** Normalized Raman intensity over the CH and OH vibration bands (2750-3800cm$^{-1}$). The blue and black curves respectively represent the Raman spectra of pure water and a weighed sum of the spectra of pure $K_{10}$ and pure $D_{10}$. The colored straight lines represent the Raman spectra of condensates of different radii (see color bar). **(c)** Phasor representation of the data in (b), the dotted line is a guide to the eye that attests of the alignment of the datapoints. The color of each data point corresponds to that of its Raman spectrum counterpart presented in (b). The dotted line is a guide to the eye connecting the water and protein spectral phasor coordinates. **(d)** Ratio of the area below the Raman curves of condensates over the CH bond vibration region (2800-3100cm$^{-1}$) noted $A_{CH}$ and OH and NH bonds vibration region (3100-3800cm$^{-1}$) noted $A_{OH\text{-}NH}$ as a function of condensate radius. The color of the data points relates to the radii of the different condensates considered and is the same as in (b-c). The function used to fit the data was of the type $y = aR^2/(R^2 + b)$, where $a$ and $b$ are fitting constants, where $a$, $b$ and $R$ are respectively two fitting constants and condensate radius. The dotted rectangle represents the set of datapoints for which the ratio $A_{CH}/A_{OH\text{-}NH}$ no longer depends on condensate radius. **(e)** Volume fractions of the different components of the dense phase of $K_{10}$-$D_{10}$ condensates. The data was truncated according to a threshold of 1% to the maximum value of the $A_{CH}/A_{OH\text{-}NH}$ ratio.

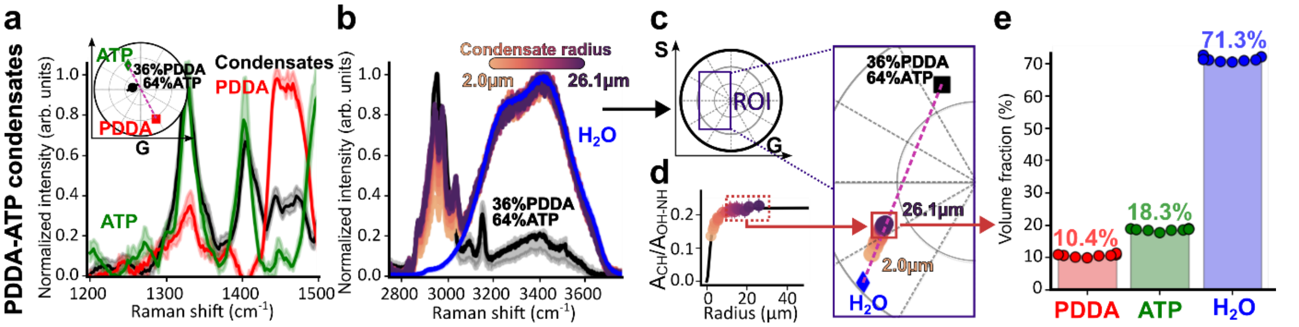


**Figure S6: Single-droplet phasor analysis of PDDA-ATP condensates. (a)** Normalized Raman intensity over the high-frequency part of the signature region. The red, green and black curves correspond to the Raman spectra of pure PDDA, pure ATP and PDDA-ATP condensates, respectively. The top left inset shows the phasor representation of the Raman spectra of pure PDDA, ATP and condensates. The proportions of PDDA and ATP were computed based on Eq. M1. **(b)** Normalized Raman intensity over the CH and OH vibration bands (2750-3800 cm$^{-1}$). The blue and black curves respectively represent the Raman spectra of pure water and a weighed sum of the spectra of pure PDDA and pure ATP. The Raman spectra of condensates are represented in color where the color bar represents condensate radius. **(c)** Phasor representation of the data in (b), the dotted line is a guide to the eye that attests of the alignment of the datapoints. The color of each data point corresponds to that of its Raman spectrum counterpart presented in (b). The dotted line is a guide to the eye connecting the water and the non-aqueous spectral phasor coordinates. **(d)** Ratio of the area below the Raman curves of condensates over the CH bond vibration region (2800-3100cm$^{-1}$) noted $A_{CH}$ and OH and NH bonds vibration region (3100-3800cm$^{-1}$) noted $A_{OH\text{-}NH}$ as a function of condensate radius. The color of the data points relates to the radii of the different condensates considered and is the same as in (b-c). The function used to fit the data was of the type $y = aR^2/(R^2 + b)$, where $a$, $b$ and $R$ are respectively two fitting constants and condensate radius. The rectangle represents the set of datapoints for which the ratio $A_{CH}/A_{OH\text{-}NH}$ no longer depends on condensate radius. **(e)** Volume fractions of the different components of the dense phase of PDDA-ATP condensates. The data was truncated according to a threshold of 1% to the maximum value of the $A_{CH}/A_{OH\text{-}NH}$ ratio.

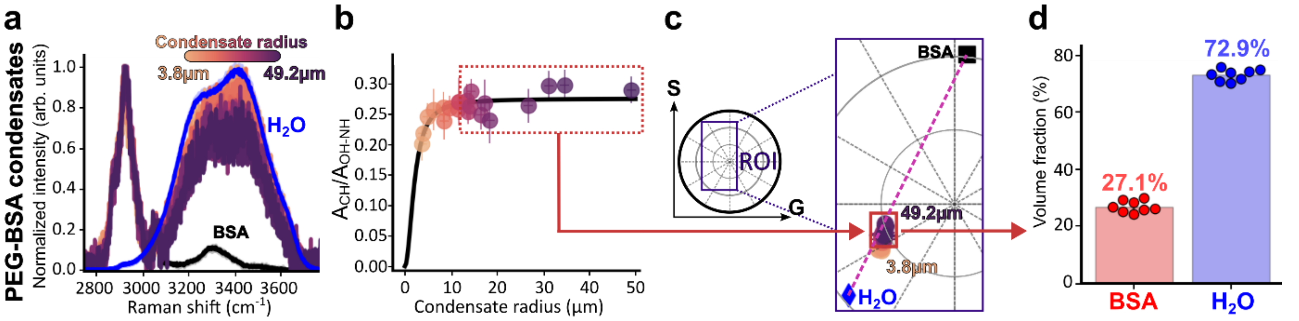


**Figure S7: Single droplet phasor analysis of BSA condensates. (a)** Normalized Raman intensity over the CH and OH vibration bands (2750-3800 cm$^{-1}$). The blue and black curves respectively represent the Raman spectra of pure water and pure BSA protein. The Raman spectra of condensates are represented in color where the color bar represents condensate radius. **(b)** Ratio of the area below the Raman curves of condensates over the CH bond vibration region (2800-3100 cm$^{-1}$) noted $A_{CH}$ and OH and NH bonds vibration region (3100-3800 cm$^{-1}$) noted $A_{OH\text{-}NH}$ as a function of condensate radius. The color of the data points relates to the radii of the different condensates considered and is the same as in (a). The dotted rectangle represents the set of datapoints for which the ratio $A_{CH}/A_{OH\text{-}NH}$ no longer depends on condensate radius. The function used to fit the data was of the type $y = aR^2/(R^2 + b)$, where $a$, $b$ and $R$ are respectively two fitting constants and condensate radius. **(c)** Phasor representation of the data in (a), the dotted line is a guide to the eye that attests of the alignment of the datapoints. The color of each data point corresponds to that of its Raman spectrum counterpart presented in (a). The dotted line is a guide to the eye connecting the water and BSA phasor coordinates. **(d)** Volume fractions of the different components of the dense phase of PEG-BSA condensates. The data was truncated according to a threshold of 1% to the maximum value of the $A_{CH}/A_{OH\text{-}NH}$ ratio.

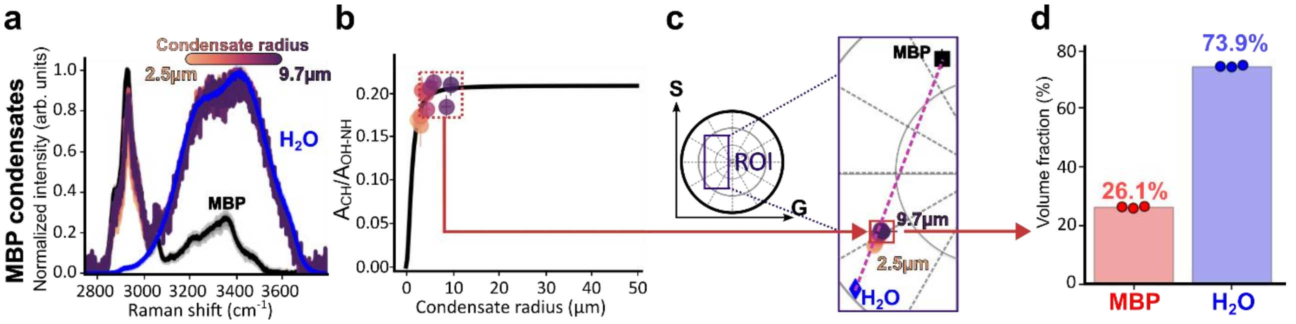


**Figure S8: Single droplet phasor analysis of MBP condensates. (a)** Normalized Raman intensity over the CH and OH vibration bands (2750-3800 $cm^{-1}$). The blue and black curves respectively represent the Raman spectra of pure water and pure MBP protein. The Raman spectra of condensates are represented in color where the color bar represents condensate radius. **(b)** Ratio of the area below the Raman curves of condensates over the CH bond vibration region (2800-3100 $cm^{-1}$) noted $A_{CH}$ and OH and NH bonds vibration region (3100-3800 $cm^{-1}$) noted $A_{OH\text{-}NH}$ as a function of condensate radius. The color of the data points relates to the radii of the different condensates considered and is the same as in (a). The dotted rectangle represents the set of datapoints for which the ratio $A_{CH}/A_{OH\text{-}NH}$ no longer depends on condensate radius. The function used to fit the data was of the type $y = aR^2/(R^2 + b)$, where $a$, $b$ and $R$ are respectively two fitting constants and condensate radius. **(c)** Phasor representation of the data in (a), the dotted line is a guide to the eye that attests of the alignment of the datapoints. The color of each data point corresponds to that of its Raman spectrum counterpart presented in (a). The dotted line is a guide to the eye connecting the water and MBP phasor coordinates. **(d)** Volume fractions of the different components of the dense phase of MBP condensates. The data was truncated according to a threshold of 1% to the maximum value of the $A_{CH}/A_{OH\text{-}NH}$ ratio.

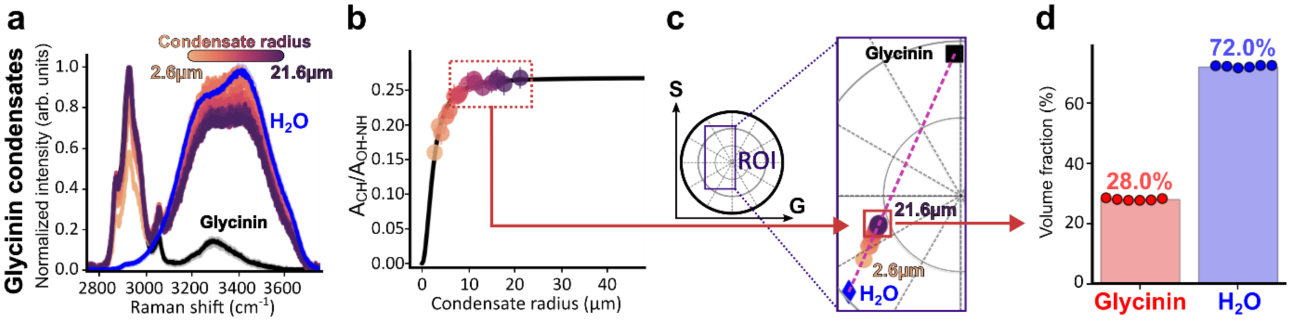


**Figure S9: Single droplet phasor analysis of glycinin condensates. (a)** Normalized Raman intensity over the CH and OH vibration bands (2750-3800 $cm^{-1}$). The blue and black curves respectively represent the Raman spectra of pure water and pure glycinin protein. The Raman spectra of condensates are represented in color where the color bar represents condensate radius. **(b)** Ratio of the area below the Raman curves of condensates over the CH bond vibration region (2800-3100 $cm^{-1}$) noted $A_{CH}$ and OH and NH bonds vibration region (3100-3800 $cm^{-1}$) noted $A_{OH\text{-}NH}$ as a function of condensate radius. The color of the data points relates to the radii of the different condensates considered and is the same as in (a). The dotted rectangle represents the set of datapoints for which the ratio $A_{CH}/A_{OH\text{-}NH}$ no longer depends on condensate radius. The function used to fit the data was of the type $y = aR^2/(R^2 + b)$, where $a$, $b$ and $R$ are respectively two fitting constants and condensate radius. **(c)** phasor representation of the data in (a), the dotted line is a guide to the eye that attests of the alignment of the datapoints. The color of each data point corresponds to that of its Raman spectrum counterpart presented in (a). The dotted line is a guide to the eye connecting the water and glycinin phasor coordinates. **(d)** Volume fractions of the different components of the dense phase of glycinin condensates. The data was truncated according to a threshold of 1% to the maximum value of the $A_{CH}/A_{OH\text{-}NH}$ ratio.

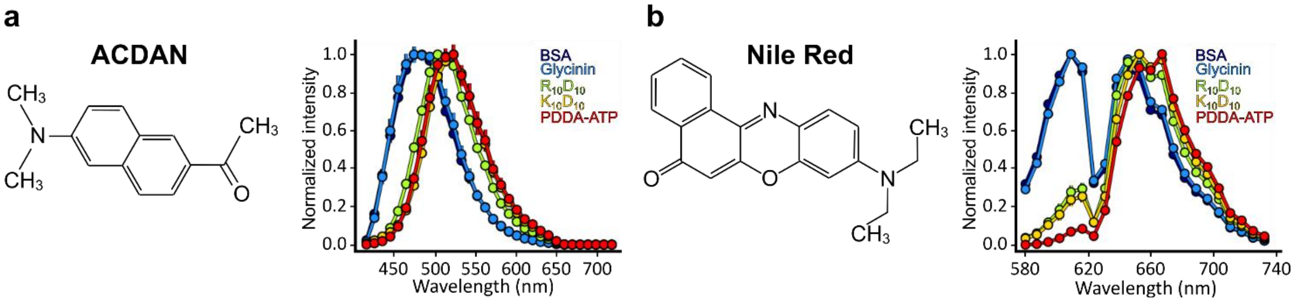


**Figure S10: ACDAN and Nile Red emission redshifts yield a similar ranking of condensate hydrophobicity. (a)** ACDAN emission spectra of the dense phase of different condensates systems introduced in Fig. 6. **(b)** Nile red emission spectra of the dense phase of different condensates systems introduced in Fig. 6.

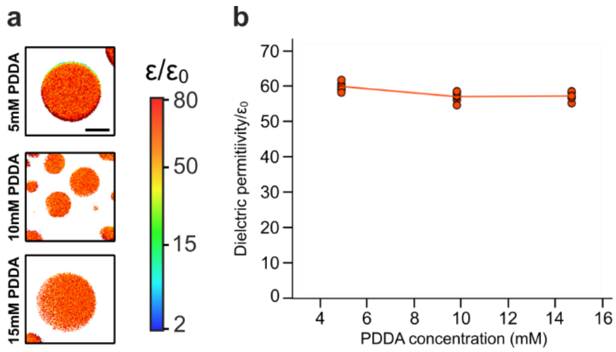


**Figure S11: Influence of PDDA concentration on the dielectric environment of biomolecular condensates. (a)** Mapping the dielectric constant of PDDA-ATP condensate systems with different total PDDA concentrations. **(b)** Quantifying the evolution of PDDA-ATP condensates as a function of PDDA concentration.

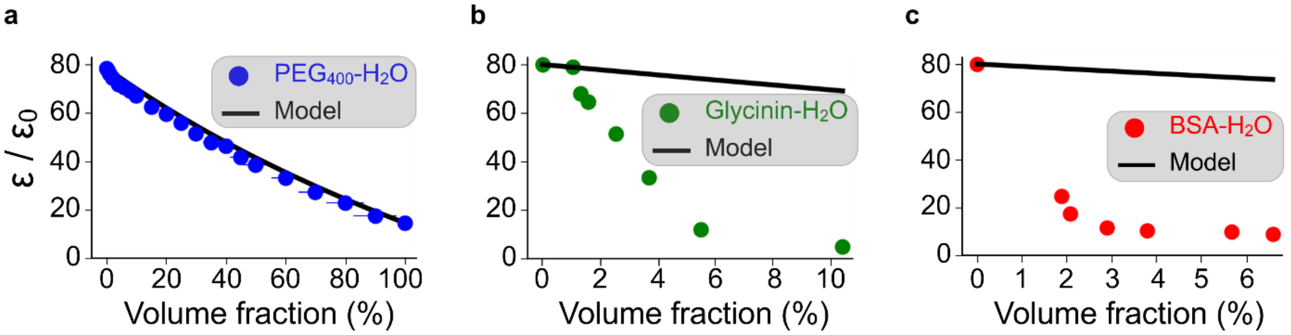


**Figure S12: ACDAN mixes unevenly in solutions containing structurally complex proteins. (a)** Comparison between dielectric constant of aqueous $PEG_{400}$ solutions measured using ACDAN fluorescence and the prediction of the Maxwell-Garnet effective medium model, a classical framework describing the effective dielectric constant of binary mixtures composed of spherical inclusions in a homogeneous solvent[1]. According to the model, the effective dielectric constant of the mixture reads $\varepsilon_{eff} = \frac{\varepsilon_{H_2O}(2\phi_{prot}(\varepsilon_{prot}-\varepsilon_{H_2O})+\varepsilon_{prot}+2\varepsilon_{H_2O})}{2\varepsilon_{H_2O}+\varepsilon_{prot}-\phi_{prot}(\varepsilon_{H_2O}-\varepsilon_{prot})}$, where $\varepsilon_{H_2O}$ and $\varepsilon_{prot}$ denote dielectric constants of pure water and pure protein respectively, and $\phi_{prot}$ is the volume fraction occupied by the protein in the mixture. The dielectric constant of aqueous $PEG_{400}$ solutions were taken from ref. [2] and used to calibrate ACDAN-based measurements of glycinin-water and BSA-water mixtures (see ref. [3] for a detailed description of the calibration method). The protein volume fractions were independently determined using Raman spectral phasor analysis (see Figs. 3 and S1). **(b-c)** Same analysis as in (a) for aqueous glycinin and BSA solutions respectively. Systematic deviations from the Maxwell-Garnett prediction indicate non-uniform mixing of ACDAN in protein solutions exhibiting structural heterogeneity.

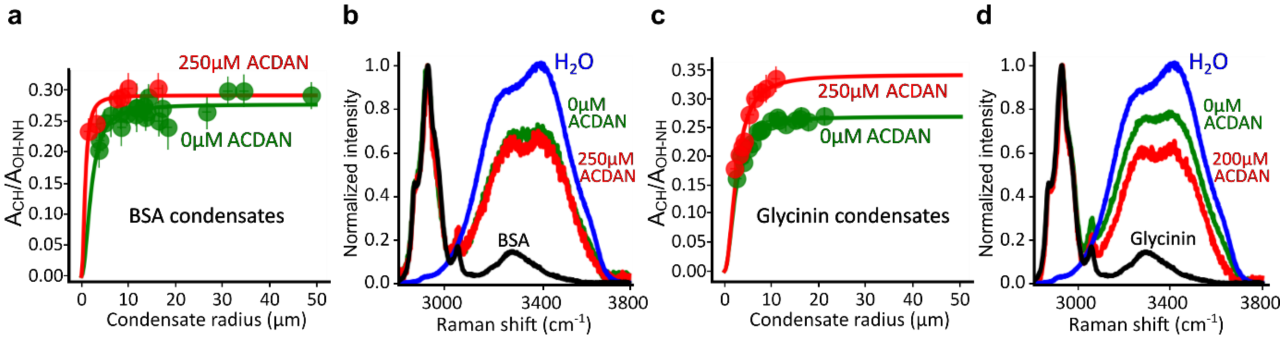


**Figure S13: Preferential ACDAN partitioning in the dense phase does not increase condensate hydration. (a)** Ratio of the area below the Raman intensity spectra of BSA condensates in the C–H stretching region (2800-3100cm$^{-1}$, $A_{CH}$) and the O–H / N–H stretching region (3100-3800cm$^{-1}$, $A_{OH-NH}$) as a function of condensate radius. The function used to fit the data was of the type $y = aR^2/(R^2 + b)$, where $a$, $b$ and $R$ are respectively two fitting constants and condensate radius. **(b)** Raman spectra of BSA condensates at varied concentrations of ACDAN. The blue, green, red and black curves respectively correspond to water, condensates without ACDAN, condensates with 250µM ACDAN, and pure BSA protein. **(c-d)** Same analysis as in (a-b) for glycinin condensates. Despite the increase in apparent dielectric constant at higher ACDAN concentrations (Fig. 6e), the systematically elevated $A_{CH}$ /$A_{OH-NH}$ ratio indicates that increased probe concentration does not enhance water partitioning into the dense phase.

**Table S1:** Summary of the specific volumes for the set of polymers used in this study.

| Protein | Parameter | Value | Unit | Reference |
|---|---|---|---|---|
| BSA | $\nu_{BSA}$ | 0.73 | L.kg$^{-1}$ | 4 |
| Glycinin | $\nu_{Glycinin}$ | 0.73 | L.kg$^{-1}$ | 5 |
| $PEG_{400}$ | $\nu_{PEG_{400}}$ | 0.89 | L.kg$^{-1}$ | 6 |
| $PEG_{3350}$ | $\nu_{PEG_{3350}}$ | 0.83 | L.kg$^{-1}$ | 7 |
| $PEG_{8000}$ | $\nu_{PEG_{8000}}$ | 0.83 | L.kg$^{-1}$ | 8 |
| Dextran | $\nu_{Dextran}$ | 0.62 | L.kg$^{-1}$ | 8 |
| Water | $\nu_{H_2O}$ | 1 | L.kg$^{-1}$ | |
